# Long-term potentiation at pyramidal cell to somatostatin interneuron synapses controls hippocampal network plasticity and memory

**DOI:** 10.1101/2021.11.23.469739

**Authors:** Azam Asgarihafshejani, Ève Honoré, François-Xavier Michon, Isabel Laplante, Jean-Claude Lacaille

## Abstract

Hippocampal somatostatin (SOM) cells are dendrite-projecting inhibitory interneurons. CA1 SOM cells receive major excitatory inputs from pyramidal cells (PC-SOM synapses) which show mGluR1a- and mTORC1-mediated long-term potentiation (LTP). PC-SOM synapse LTP contributes to CA1 network metaplasticity and memory consolidation, but whether it is sufficient to regulate these processes remains unknown. Here we used optogenetic stimulation of CA1 pyramidal cells and whole cell recordings in slices to show that optogenetic theta burst stimulation (TBS_opto_) produces LTP at PC-SOM synapses. At the network level, we found that TBS_opto_ differentially regulates metaplasticity of pyramidal cell inputs: enhancing LTP at Schaffer collateral synapses and depressing LTP at temporo-ammonic synapses. At the behavioral level, we uncovered that in vivo TBS_opto_ regulates learning-induced LTP at PC-SOM synapses, as well as contextual fear memory. Thus, LTP of PC-SOM synapses is a long-term feedback mechanism controlling pyramidal cell synaptic plasticity, sufficient to regulate memory consolidation.

## INTRODUCTION

Hippocampal interneurons are heterogeneous populations of GABAergic inhibitory cells with varied morphological, molecular and electrophysiological properties, as well as specialized network functions (Pelkey et al., 2017, Klausberger and Somogyi, 2008, Booker and Vida, 2018, Freund and Buzsaki, 1996, Bezaire and Soltesz, 2013). CA1 somatostatin-expressing (SOM) cells are a major interneuron subgroup comprised notably of Oriens-Lacunosum/Moleculare (O-LM) cells, bistratified cells and long-range projecting cells (Katona et al., 2014, Pelkey et al., 2017, Honore et al., 2021). SOM cells have axons that target dendrites of pyramidal cells (Muller and Remy, 2014, Lovett-Barron et al., 2014, Klausberger et al., 2003, Pelkey et al., 2017) as well as other interneurons (Fuhrmann et al., 2015, Leao et al., 2012). SOM cells suppress spike rate and burst firing of pyramidal cells in vitro (Lovett-Barron et al., 2012) and of place cells during exploration in vivo (Royer et al., 2012). SOM cells fire with specific phase-coupling during hippocampal theta and gamma network oscillations that are hallmarks of hippocampal function and occur during spatial navigation, memory tasks, and rapid-eye-movement sleep (Klausberger and Somogyi, 2008, Katona et al., 2014). Indeed, SOM cell regulation of pyramidal neuron activity is critical for hippocampus-dependent learning and memory, as silencing SOM cells during contextual fear conditioning impairs long-term contextual memory (Lovett-Barron et al., 2014).

SOM cells, like other hippocampal interneurons, show long-term synaptic plasticity of their input synapses (Kullmann et al., 2012, Pelletier and Lacaille, 2008, Honore et al., 2021). CA1 pyramidal cell excitatory synapses onto SOM cells (PC-SOM synapses) display a Hebbian form of long-term potentiation (LTP) (Vasuta et al., 2015, Perez et al., 2001, Lapointe et al., 2004, Honore et al., 2021, Croce et al., 2010) dependent on metabotropic glutamate receptor subtype 1a (mGluR1a) activation and postsynaptic Ca^2+^ rise (Lapointe et al., 2004, Perez et al., 2001). This form of long-term plasticity is cell-specific and absent in parvalbumin-expressing cells (Vasuta et al., 2015) and radiatum interneurons (Perez et al., 2001). LTP at input synapses enables long-term regulation of interneuron firing (Croce et al., 2010) and pyramidal cell inhibition (Lapointe et al., 2004).

A critical SOM cell function is the differential regulation of afferents onto pyramidal cell dendrites: via distal dendritic inhibition, SOM cells downregulate activity and LTP in the temporo-ammonic pathway (TA-PC); whereas via proximal dendritic disinhibition (inhibition of other inhibitory interneurons), they upregulate activity and LTP in the CA3-CA1 Schaffer collateral pathway (SC-PC) (Leao et al., 2012). Remarkably, Hebbian LTP at PC-SOM synapses increases subsequent LTP at SC-PC synapses (Vasuta et al., 2015) and decreases subsequent LTP at TA-PC synapses (Sharma et al., 2020), showing that plasticity at PC-SOM input synapses controls durably CA1 network metaplasticity (Honore et al., 2021).

LTP at PC-SOM synapses can be persistent, as repeated activation of mGluR1a induces LTP of excitatory synapses lasting hours and involving gene transcription and mRNA translation (Ran et al., 2009, Artinian et al., 2019). mTORC1 signaling controls translation during LTP at PC-SOM synapses (Ran et al., 2009, Artinian et al., 2019). Interestingly, mTORC1 activity in SOM cells is required for learning-induced LTP of PC-SOM synapses, regulation of CA1 network plasticity, and long-term consolidation of spatial and contextual fear memory (Artinian et al., 2019). However, whether LTP at PC-SOM synapses is sufficient for the regulation of CA1 network metaplasticity and hippocampus-dependent memory remains to be determined.

In the present study, we address this question by using optogenetics to induce long-term plasticity at PC-SOM excitatory synapses. We found that optogenetic stimulation of pyramidal cells induces LTP at PC-SOM synapses in slices. Furthermore, optogenetic induction of LTP at PC-SOM synapses bi-directionally controls CA1 network plasticity by facilitating SC-PC LTP and suppressing TA-PC LTP. Moreover, we uncover that optogenetic induction of LTP at PC-SOM synapses *in vivo* regulates learning-induced LTP at these synapses and controls contextual fear memory. Our findings suggest that LTP at PC-SOM synapses is sufficient to control the state of plasticity in the CA1 network and the consolidation of contextual fear memory, uncovering a long-term feedback control mechanism via PC-SOM synapses sufficient for regulation of CA1 pyramidal cell metaplasticity and hippocampus-dependent memory.

## RESULTS

### Optogenetically-induced LTP at PC-SOM synapses

CA1 pyramidal cells are the major excitatory synaptic inputs to SOM interneurons in CA1 stratum oriens (Croce et al., 2010, Honore et al., 2021). We used a combination of optogenetic stimulation after injection of AAV2/9-CaMKIIa-hChR2(E123T/T159C)-mCherry in SOM-Cre-EYFP mice to activate CA1 pyramidal cells, and of whole cell current clamp recordings in slices with synaptic inhibition intact to monitor excitatory postsynaptic potentials (EPSPs) in CA1 SOM interneurons resulting from activation of PC-SOM synapses (Figure 1A). Local optogenetic stimulation (470 nm) through the objective was adjusted (pulse duration 0.5-2.0 ms) to elicit light-evoked EPSPs in SOM interneurons (Figures 1B-G and S1A-G), similar to EPSPs evoked by electrical stimulation (Figure S2B-F) (Vasuta et al., 2015). Theta burst stimulation (TBS) is an effective induction protocol for eliciting Hebbian LTP at PC-SOM excitatory synapses (Perez et al., 2001, Croce et al., 2010, Vasuta et al., 2015). Therefore, we developed an optogenetic TBS protocol (TBS_opto_; bursts of 4 light pulses at 80 Hz, given 5 times with 300 ms inter-burst intervals, repeated 3 times at 30 s intervals) effective in eliciting EPSP summation and burst firing of SOM cells (Figure 1B). The TBS_opto_ protocol induced LTP of light-evoked EPSPs at PC-SOM synapses. Light-evoked EPSP amplitude showed a slow onset, gradual increase, lasting at least 30 min after TBS_opto_ (132.0% of baseline at 20-30 min post-induction; Figure 1B and 1J).

**Figure 1.**
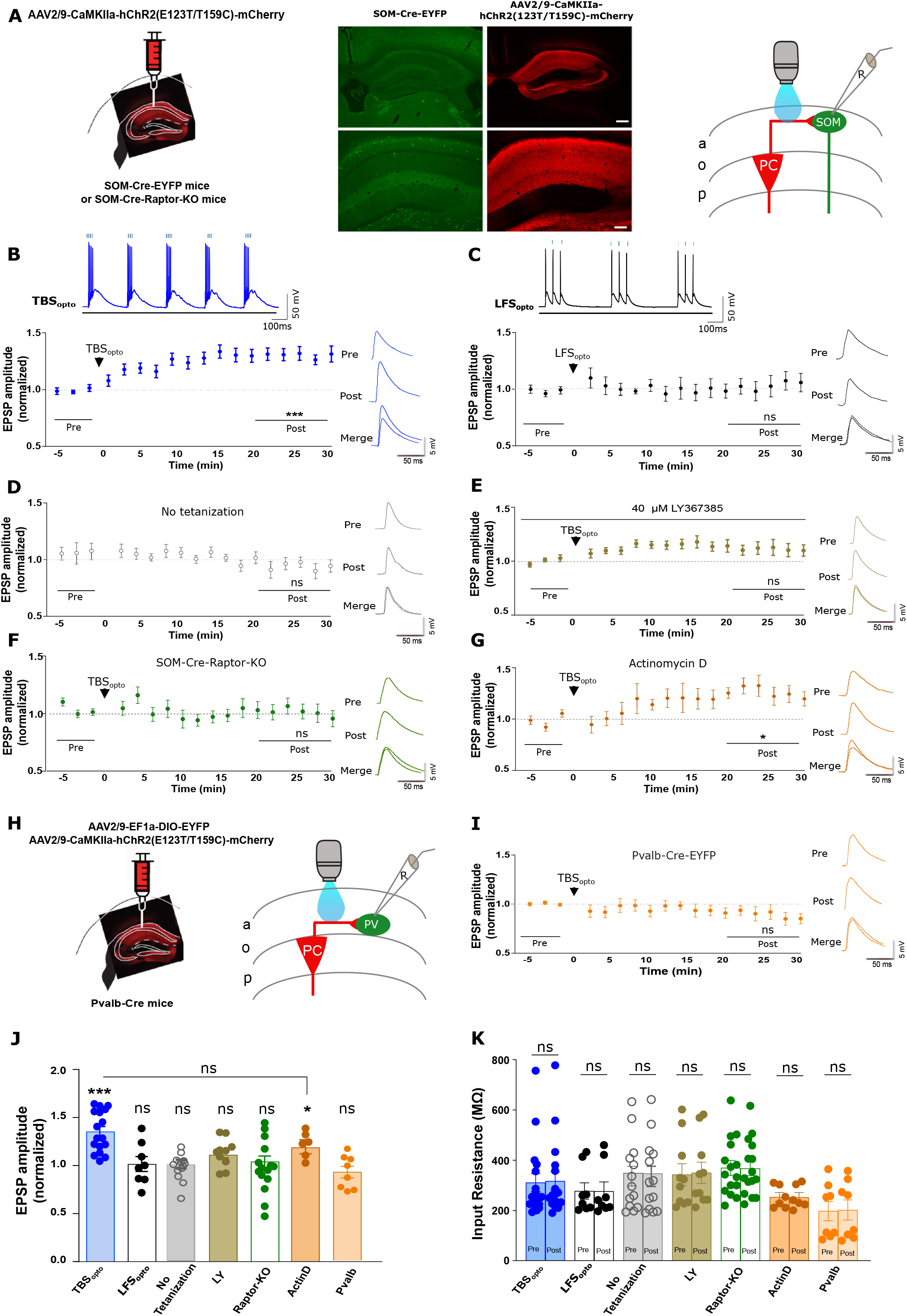
Optogenetically-induced LTP at PC-SOM synapses. (A) Left: schematic of experimental paradigm showing viral injections in dorsal hippocampus of SOM-Cre-EYFP or SOM-Cre-Raptor-KO mice. Middle: representative images of EYFP expression in SOM cells and mCherry expression in CA1 pyramidal cells (calibration bar: 100 μm bottom; 250 μm top). Right: diagram of local optogenetic stimulation of CA1 pyramidal cells and whole cell recording from SOM interneurons in stratum oriens. (B and C) Bottom left: time plots of light-evoked EPSP amplitude for all SOM cells, showing LTP at PC-SOM synapses following TBS_opto_ (B; n = 17 cells) but not after LFS_opto_ (C; n = 8 cells). Right: example of representative cells of average EPSPs before (Pre) and 20-30 min after (Post) TBS_opto_ or LFS_opto_. Top: EPSP summation and cell firing during TBS_opto_ (80 Hz) or LFS_opto_ (20 Hz). Paired t-tests; *** *p* < 0.0001; ns, *p* > 0.05. See Figure S1A and S1B for the respective time plots of EPSP amplitude of representative cells in (B) and (C), respectively. (D - G) Left: time plots of light-evoked EPSP amplitude for all SOM cells, showing absence of LTP at PC-SOM synapses following no tetanization (D; n = 13 cells), TBS_opto_ in the presence of the mGluR1a antagonist LY367385 (40 μM) (E; n = 10 cells), or TBS_opto_ in SOM-Cre-Raptor-KO mice (F; n = 15 cells); but LTP after TBS_opto_ in slices pre-incubated with the transcription inhibitor actinomycin D (2μM) (G; n = 6 cells). Right: representative example of average EPSPs before (Pre) and after (Post) each treatment condition. Paired t-tests; * *p* < 0.05; ns, *p* > 0.05. See Figure S1C, S1D and S1F for the respective time plots of EPSP amplitude of representative cells in (D), (E), (F) and (G), respectively; and Figure S1G for representative cell and S1H for time plot for all cells, showing LTP after TBS_opto_ in DMSO (vehicle for actinomycin D). (H) Schematic of experimental paradigm with (left) viral injections in dorsal hippocampus of Pvalb-Cre mice and (right) local optogenetic stimulation of pyramidal cells and recording from CA1 PV interneurons. (I) Time plot of light-evoked EPSP amplitude for all PV cells (left) and example of average EPSPs from a representative cell (right), showing absence of LTP at PC-PV synapses following TBS_opto_ (n = 8 cells). Paired t-test; ns, *p* > 0.05. See Figure S1I for the time plot of EPSP amplitude of the representative PV cell in (I). (J) Summary graph of EPSP amplitude for all cells at 20-30 min post induction, showing LTP after TBS_opto_ in ACSF and after incubation with actinomycin D, but no LTP after LFS_opto_, in absence of tetanization, or after TBS_opto_ in LY367385, in Som-Cre-RAPTOR mice, or in PV cells. Paired t-tests (Post vs Pre in B-I) and unpaired t-test (Post in B vs Post in G); *** p < 0.0001; * p < 0.05; ns, *p* > 0.05. (K) Summary graph for all cells, showing no change in cell input resistance at 20-30 min post induction in any experimental group. Wilcoxon matched pairs signed rank test (TBS_opto_ and LFS_opto_), or paired t-test (other groups); ns, *p* > 0.05. Data are represented as mean± SEM here and below. *p < 0.05, **p < 0.01, ***p < 0.001; p > 0.5 is considered not significant (ns). Statistical tests and actual p values for each comparison here and below are given in Table S1.

Next, we examined the mechanisms involved in the optogenetically-induced LTP at PC-SOM synapses. First, low frequency optogenetic stimulation (LFS_opto_; 20 Hz) that elicited EPSP summation but a lower rate of firing in SOM cells (Figure 1C) did not induce lasting changes in light-evoked EPSP amplitude (103.0% of baseline at 20-30 min post-induction; Figure 1C and 1J, Figure S1B). Second, in recordings with no optogenetic tetanization, light-evoked EPSP amplitude was unchanged over the same time period (95.7% of baseline at 20-30 min post-induction; Figure 1D and 1J, Figure S1C). Third, to determine whether the optogenetically-induced LTP at PC-SOM synapses was mGluR1a-dependent, TBS_opto_ was given in the presence of the mGluR1a antagonist LY367385 (40 μM) (Figure 1E). In these conditions, TBS_opto_ failed to induce lasting changes in EPSP amplitude (111.0% of baseline at 20-30 min post-induction; Figure 1E and 1J, Figure S1D). Fourth, to determine whether optogenetically-induced LTP at PC-SOM synapses is dependent on mTORC1-mediated translation, we used SOM-Raptor-KO mice that are deficient in mTORC1 signaling and late-LTP in SOM cells (Artinian et al., 2019). In these mice, TBS_opto_ failed to induce lasting changes in light-evoked EPSP amplitude (104.0% of baseline at 20-30 min post-induction; Figure 1F and 1J, Figure S1E). Fifth, we examined if optogenetically-induced LTP requires transcription by pre-incubating hippocampal slices with 2 μM of the irreversible transcription inhibitor actinomycin D for 15 min prior to recording (Yuan and Burrell, 2013, Younts et al., 2016). In these conditions, TBS_opto_ induced a slow onset LTP of light-evoked EPSP amplitude (119.5% of baseline at 20-30 min post-induction; Figure 1G and 1J, Figure S1F) that was comparable to the LTP induced in control (Figure 1B and 1J) or in vehicle (DMSO; Figure S1G-H). Sixth, we determined if optogenetic stimulation elicited LTP at PC synapses onto parvalbumin-expressing interneurons (PC-Pvalb synapses) using injection of AAV2/9-CaMKIIa-hChR2(E123T/T159C)-mCherry and AAV2/9-EF1a-DIO-EYFP in Pvalb-Cre mice (Figure 1H). In these mice, TBS_opto_ failed to induce lasting increases in light-evoked EPSPs at PC-Pvalb synapses (90.0% of baseline at 20-30 min post-induction; Figure 1I-J, Figure S1I). In all these experiments, cell input resistance was unchanged after TBS_opto_ (Figure 1K). Together, these results suggest that optogenetic stimulation of CA1 pyramidal cells produces a frequency-dependent LTP at PC-SOM synapses that is dependent on mGluR1a activation and mTORC1-mediated translation, but independent of transcription. Moreover, optogenetic stimulation of CA1 pyramidal cells does not induce LTP at PC-Pvalb synapses.

These properties of optogenetically-induced LTP at PC-SOM synapses are similar to those previously reported for Hebbian LTP at excitatory synapses onto SOM cells characterized using electrical stimulation (Perez et al., 2001, Croce et al., 2010, Honore et al., 2021). So, next we determined if TBS_opto_ can induce LTP of EPSPs evoked in SOM cells by electrical stimulation (Figure S2A). We found that TBS_opto_ induced a similar LTP of electrically-evoked EPSPs that was dependent on mGluR1a and mTORC1, required tetanization and was absent in mice that received only the control CaMKIIa-mCherry injection without hChR2 (Figure S2B-G), indicating similar mechanisms as at PC-SOM synapses activated optogenetically.

### TBS_opto_ differentially regulates LTP at Schaffer collateral and temporo-ammonic synapses

CA1 SOM cells (notably OLM cells) differentially control transmission at Schaffer collateral and temporo-ammonic synapses of CA1 pyramidal cells (Leao et al., 2012). In consequence, Hebbian LTP at excitatory synapses onto SOM cells is associated with increased LTP at Schaffer collateral inputs and reduced LTP at temporo-ammonic synapses of CA1 pyramidal cells (Vasuta et al., 2015, Artinian et al., 2019, Sharma et al., 2020). So, next we studied if TBS_opto_-induced LTP at PC-SOM synapses was sufficient to regulate metaplasticity of the CA1 network using whole-field optogenetic stimulation and field potential recordings in slices with synaptic inhibition intact. First, we established that whole-field optogenetic stimulation also elicited LTP at PC-SOM synapses (Figure 2A). Whole-field TBS_opto_ elicited EPSP summation and burst firing of SOM cells, and induced LTP of light-evoked EPSP amplitude (125.0% of baseline at 20-30 min post-induction; Figure 2B and S3E). LTP was absent in cells that received no optogenetic tetanization (light-evoked EPSP amplitude 97.8% of baseline at 30 min post-induction; Figure S3B and S3E), or with TBS_opto_ given in the presence of the mGluR1a antagonist LY367385 (40 μM) (light-evoked EPSP amplitude 97.0% of baseline at 20-30 min post-induction; Figure S3C and S3E), or in SOM cells of SOM-Raptor-KO mice (light-evoked EPSP amplitude 96.3% of baseline at 20-30 min post-induction; Figure S3D and S3E). Thus, whole-field TBS_opto_ induced a similar mGluR1a- and mTORC1-mediated LTP at PC-SOM synapses as local optogenetic stimulation.

**Figure 2.**
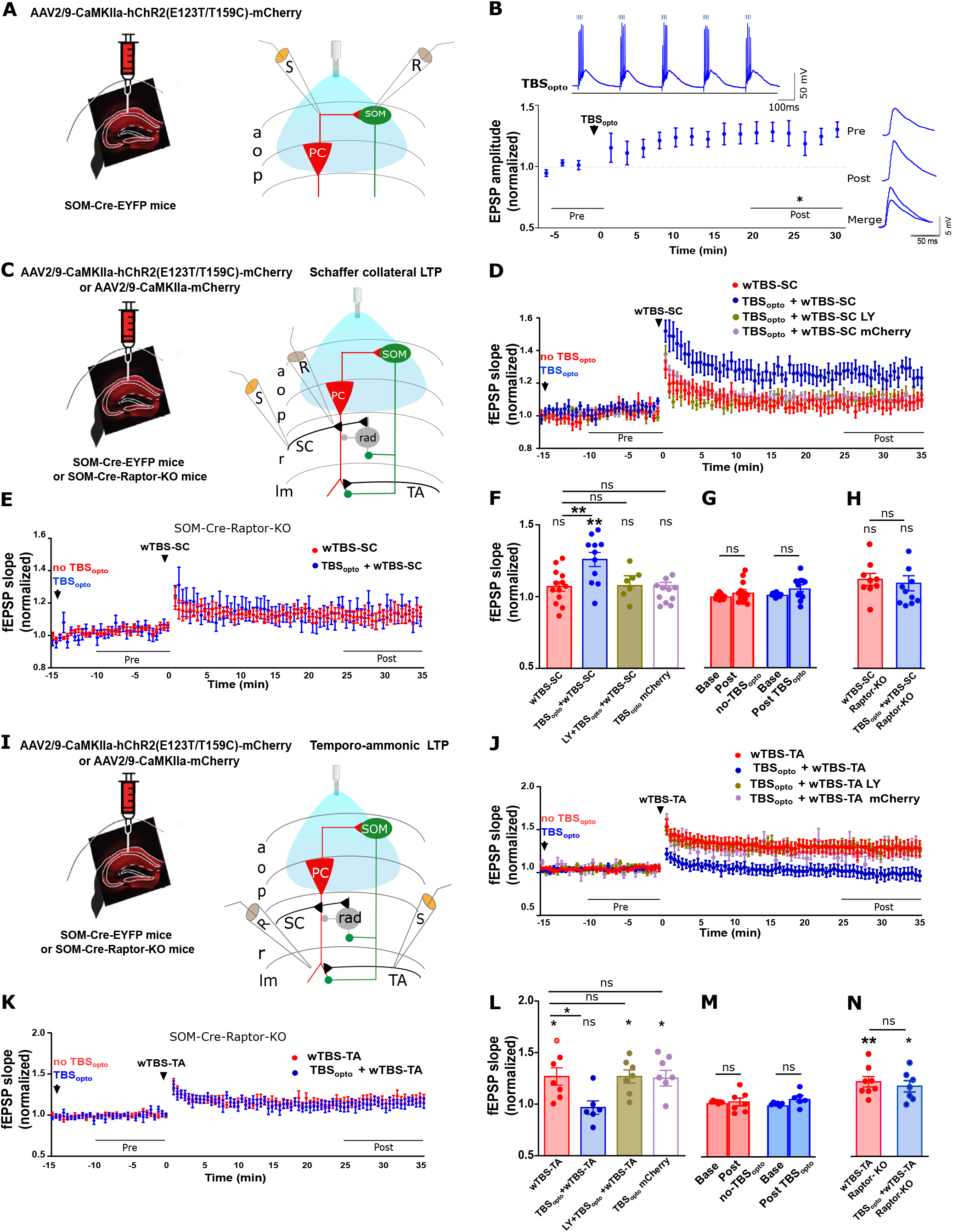
Whole-field TBS_opto_ induces LTP at PC-SOM synapses and differentially regulates LTP at SC-PC and TA-PC synapses. (A) Left: schematic of experimental paradigm with hippocampal viral injections in SOM-Cre-EYFP mice. Right: diagram of whole-field optogenetic stimulation of CA1 pyramidal cells and whole cell recording from SOM interneurons. (B) Left bottom: time plot of light-evoked EPSP amplitude for all SOM cells, showing LTP at PC-SOM synapses following TBS_opto_ (n = 7 cells). Left top: EPSP summation and cell firing during TBS_opto_ in a representative cell. Right: example from a representative cell of average light-evoked EPSPs before (Pre) and 20-30 min after (Post) TBS_opto_. Paired t-test; * *p* < 0.05. See Figure S3A to S3E for the dependence of whole-field TBS_opto_-induced LTP on tetanization (S3B and S3E), mGluR1a (S3C and S3E) and mTORC1 (S3D and S3E). (C) Left: schematic of experimental paradigm with hippocampal viral injections in SOM-Cre-EYFP or SOM-Cre-Raptor-KO mice. Right: diagram of whole-field optogenetic stimulation of CA1 pyramidal cells, with electrical stimulation and recording of SC-PC fEPSPs in stratum radiatum. (D) Time plot of fEPSP slope for all slices, showing that LTP at SC-PC synapses (n = 13 slices, red) is facilitated by prior application of TBS_opto_ (n = 11 slices, blue), but not in the presence of the mGluR1a antagonist LY367385 (n = 7 slices, brown) nor in slices from mice with control CaMKIIa-mCherry injection without hChR2 (n = 11 slices, magenta). See Figure S3G and S3H for time plots of fEPSP slope in representative slices without and with TBS_opto_, respectively. (E) Time plot of fEPSP slope for all slices from SOM-Cre-Raptor-KO mice, showing that prior application of TBS_opto_ (n = 10 slices, blue) does not facilitate LTP at SC-PC synapses (n = 9 slices; red) in these mutant mice. (F) Summary graph of fEPSP slope for all slices at 25-35 min post induction of LTP at SC-PC synapses, showing that LTP at SC-PC synapses (red) is facilitated by prior application of TBS_opto_ (blue), but not in LY367385 (brown) or in slices from mice injected with mCherry (magenta). Paired t-tests (Pre vs Post); ANOVA, Tukey’s multiple comparisons tests (Post wTBS-SC vs Post other conditions); * p < 0.05, ns *p* > 0.05. (G) Summary bar graph of fEPSP slope before (−10 to 0 min) and after (15-25 min) TBS_opto_, showing that basal transmission at SC-PC synapses is unchanged after TBS_opto_ or no TBS_opto_. Paired t-test, ns *p* > 0.05. (G) Summary graph of fEPSP slope for all slices at 25-35 min post induction of LTP at SC-PC synapses, showing that LTP at SC-PC synapses (red) is not facilitated by prior application of TBS_opto_ (blue) in SOM-Cre-Raptor-KO mice. Unpaired t-test, ns *p* > 0.05. (I) Left: schematic of experimental paradigm with hippocampal viral injections in SOM-Cre-EYFP or SOM-Cre-Raptor-KO mice. Right: diagram of whole-field optogenetic stimulation of CA1 pyramidal cells, with electrical stimulation and recording of TA-PC fEPSPs in stratum lacunosum-moleculare. (J) Time plot of fEPSP slope for all slices, showing that LTP at TA-PC synapses (n = 7 slices, red) is depressed by prior application of TBS_opto_ (n = 6 slices, blue), but not by TBS_opto_ in the presence of the mGluR1a antagonist LY367385 (n = 7 slices, brown), nor by TBS_opto_ in slices from mice with CaMKIIa-mCherry injection (n = 7 slices, magenta). See Figure S3J and S3K for time plots of fEPSP slope in representative slices without and with TBS_opto_, respectively. (K) Time plot of fEPSP slope for all slices from SOM-Cre-Raptor-KO mice, showing that LTP at TA-PC synapses (n = 8 slices; red) is not depressed by prior application of TBS_opto_ (n = 7 slices, blue) in these mutant mice. (L) Summary graph of fEPSP slope for all slices at 25-35 min post induction of LTP at TA-PC synapses, showing LTP at TA-PC synapses (red) is depressed by prior application of TBS_opto_ (blue), but not by TBS_opto_ in LY367385 (brown) or in slices from mice injected with mCherry (magenta). Paired t-tests (Pre vs Post); ANOVA, Tukey’s multiple comparisons tests (Post wTBS-SC vs Post other conditions); * p < 0.05, ns *p* > 0.05. (M) Summary bar graph of fEPSP slope before (−10 to 0 min) and after (15-25 min) TBS_opto_, showing that basal transmission at TA-PC synapses is unchanged after TBS_opto_ or no TBS_opto_. Paired t-test, ns *p* > 0.05. (N) Summary graph of fEPSP slope for all slices at 25-35 min post induction of LTP at TA-PC synapses, showing that LTP at TA-PC synapses (red) is not depressed by prior application of TBS_opto_ (blue) in SOM-Cre-Raptor-KO mice. Paired t-tests (Pre vs Post); unpaired t-test (wTBS-TA vs TBS_opto_ +wTBS-TA); ** p < 0.01, * p < 0.05, ns *p* > 0.05.

Next, we examined the long-lasting effects of whole-field TBS_opto_ on plasticity at Schaffer collateral synapses (SC-PC synapses) and temporo-ammonic synapses (TA-PC synapses) onto CA1 PCs in slices with synaptic inhibition intact. In these experiments, we used whole-field optogenetic stimulation and fEPSP recordings so that we could obtain long-term stable recordings of synaptic responses, for a 15 min baseline period before TBS_opto_ and a 25 min period after TBS_opto_, to assess effects of TBS_opto_ on basal transmission, and for another 35 min period after tetanization of SC-PC or TA-PC synapses, to assess effects on LTP at these synapses (Figure 2). First, we investigated regulation of plasticity at SC-PC synapses (Figure 2C and Figure S3F). A weak electrical TBS (wTBS) of the SC pathway failed to induce LTP of fEPSP slope (107.0% of baseline at 25-35 min post-induction; Figure 2D and Figure S3G). In contrast, when the wTBS was preceded 25 min earlier by TBS_opto_, LTP was induced at SC-PC synapses (125.7% of baseline at 25-35 min post-induction; Figure 2D and 2F, and Figure S3H). Thus, TBS_opto_ facilitated LTP at SC-PC synapses. TBS_opto_ did not affect basal transmission at SC-PC synapses (Figure 2G). In mice with control CaMKIIa-mCherry injection without hChR2, TBS_opto_ failed to facilitate LTP at SC-PC synapses (104.0% of baseline at 25-35 min post-induction; Figure 2D and 2F), indicating that the LTP facilitation was not due to unspecific effects of light stimulation. To determine if the facilitation of LTP at SC-PC synapses was due to plasticity at SOM-PC synapses induced by TBS_opto_, we examined if the facilitation was dependent on mGluR1a and on mTORC1 in SOM cells. When wTBS was preceded 25 min earlier by TBS_opto_ in the presence of the mGluR1a antagonist LY367385 (40 μM), the facilitation of LTP at SC-PC synapses was absent (107.0% of baseline at 25-35 min post-induction; Figure 2D and 2F). Similarly, when wTBS was preceded 25 min earlier by TBS_opto_ in slices from SOM-Raptor-KO mice, the facilitation of LTP at SC-PC synapses was also absent (112.0% of baseline at 25-35 min post-induction without TBS_opto_ and 1007.0% of baseline at 25-35 min post-induction with TBS_opto_; Figure 2E and 2H). These results suggests that the facilitation of LTP at SC-PC synapses was due to mGluR1a- and mTORC1-dependent plasticity at SOM-PC synapses induced by TBS_opto_.

Second, we investigated regulation of plasticity at TA-PC synapses (Figure 2I and Figure S3I). A weak electrical TBS (wTBS) of the TA pathway induced LTP of fEPSP slope (126.7% of baseline at 25-35 min post-induction; Figure 2J and 2L and Figure S3J). In contrast, when the wTBS was preceded 25 min earlier by TBS_opto_, LTP was prevented at TA-PC synapses (95.5% of baseline at 25-35 min post-induction; Figure 2J and 2L, and Figure S3K). Thus, TBS_opto_ depressed LTP at TA-PC synapses. TBS_opto_ did not affect basal transmission at TA-PC synapses (Figure 2M). In mice with control CaMKIIa-mCherry injection without hChR2, TBS_opto_ failed to depress LTP at TA-PC synapses (127.0% of baseline at 25-35 min post-induction; Figure 2J and 2L), indicating that the LTP depression was not due to unspecific effects of light stimulation. The depression of LTP at TA-PC synapses was likely due to plasticity at SOM-PC synapses induced by TBS_opto_ since it was dependent on mGluR1a and on mTORC1 in SOM cells. The depression of LTP at TA-PC synapses was absent when wTBS was preceded by TBS_opto_ in the presence of the mGluR1a antagonist LY367385 (126.0% of baseline at 25-35 min post-induction; Figure 2J and 2L). Similarly, the depression of LTP at TA-PC was absent when wTBS was preceded by TBS_opto_ in slices from SOM-Raptor-KO mice (123.0% of baseline at 25-35 min post-induction without TBS_opto_ and 118.9% of baseline at 25-35 min post-induction with TBS_opto_; Figure 2K and 2N). These results suggest that the TBS_opto_-induced depression of LTP at TA-PC synapses is the result of mGluR1a- and mTORC1-dependent plasticity at SOM-PC synapses. Thus, TBS_opto_-induced LTP at PC-SOM synapses appears sufficient to regulate metaplasticity of the CA1 network, upregulating LTP at SC-PC synapses and downregulating LTP at TA-PC synapses.

### TBS_opto_ interferes with contextual fear memory consolidation

At the behavioral level, mTORC1 activity in SOM cells, that is required for learning-induced LTP of PC-SOM synapses, contributes to long-term consolidation of spatial and contextual fear memory (Artinian et al., 2019). So, next we examined if LTP at PC-SOM synapses is sufficient to regulate hippocampus-dependent memory. First, we verified that, as previously reported (Lovett-Barron et al., 2014), activity of CA1 SOM cells is necessary for long-term contextual fear memory using optogenetic silencing of CA1 SOM cells with archaerhodopsin (Arch) (Vasuta et al., 2015) during contextual fear conditioning (Figure 3A and 3B). Optogenetic activation of Arch in phase with the presentation of shocks during conditioning, resulted in reduced freezing during the long-term contextual fear memory test, relative to light stimulation in absence of Arch (51.4% reduction; Figure 3C) or to activation of Arch out of phase (shifted) with presentation of shocks (64.9% reduction; Figure 3C). In the open field test (Figure S4A), silencing of CA1 SOM cells with light stimulation of Arch did not affect anxiety level (time spent in periphery or center, and ratio of time in center/periphery; Figure S4B) or locomotion (total distance traveled and zone transitions; Figure S4C) relative to mice receiving light stimulation without Arch. These results indicate that silencing SOM cells during contextual fear conditioning impairs long-term contextual fear memory, confirming that SOM cell activity is necessary during long-term contextual fear learning (Lovett-Barron et al., 2014).

**Figure 3.**
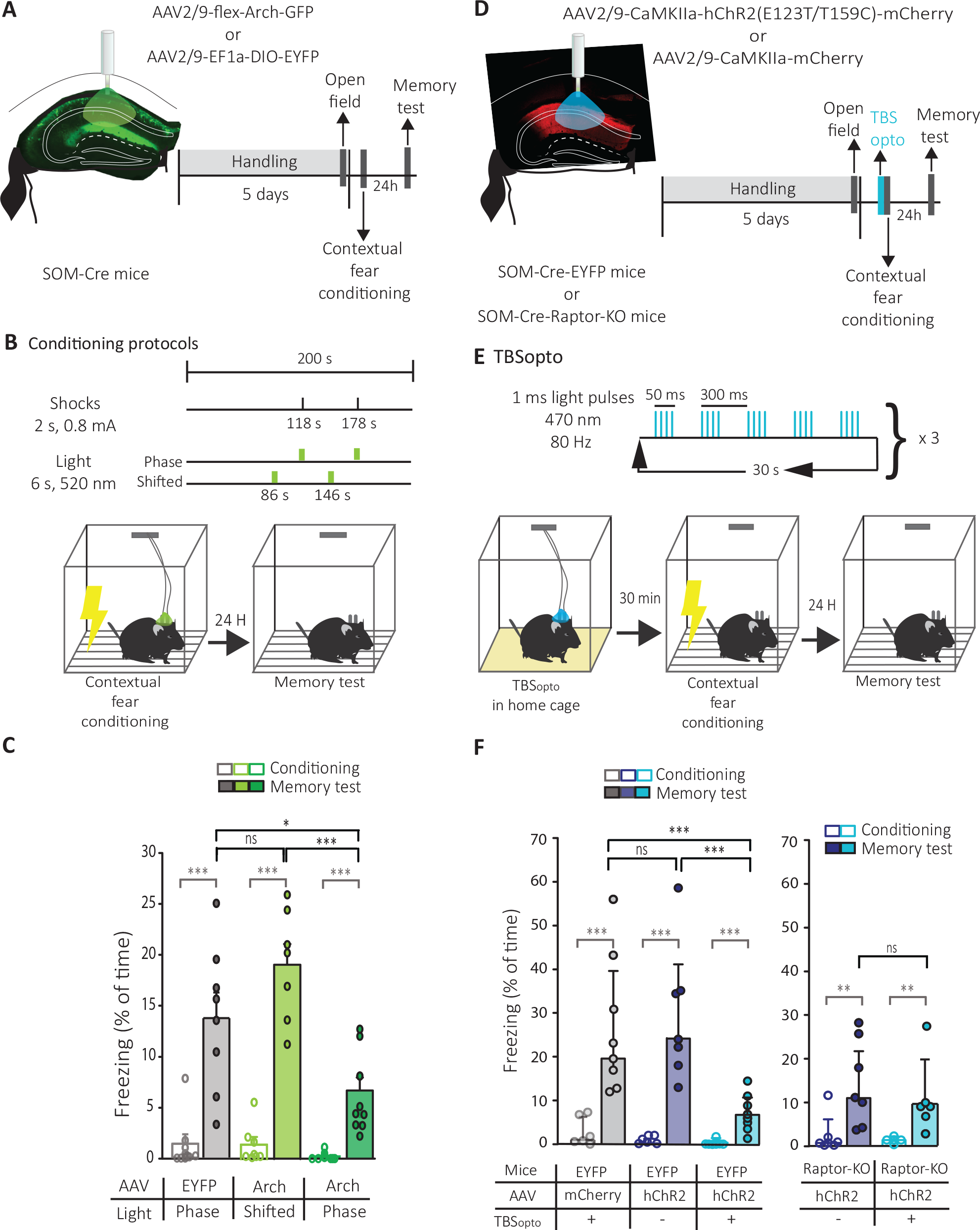
Silencing of SOM cells and TBS_opto_ impair contextual fear memory. (A) Left: schematic of cannulation and injection site of AAV2/9-flex-Arch-GFP or AAV2/9-EF1a-DIO-EYFP in dorsal CA1 hippocampus of SOM-Cre mice with representative image of Arch expression. Right: diagram of behavioral testing sequence (open field and contextual fear conditioning). (B) Experimental protocol of optogenetic stimulation (in phase or shifted with respect to shocks) during contextual fear conditioning. (C) Summary graph for all mice, showing reduced freezing in the long-term memory test in mice that received Arch activation in phase with shocks (n=9 mice, dark green), relative to those that received light stimulation without Arch (n=8 mice, gray) or Arch activation out of phase with shocks (shifted; n=7 mice, light green). Mann-Whitney Rank Sum test or Paired t-test (Pre vs Post conditioning), *** p < 0.001 (gray). One way ANOVA (Memory tests), Holm-Sidak pair-wise comparisons, * p<0.05, *** p<0.001, ns p>0.05. (D) Schematic of cannulation and injection site of AAV2/9-CaMKIIa-hChR2(E123T/T159C)-mCherry or AAV2/9-CaMKIIa-mCherry in dorsal CA1 hippocampus of SOM-Cre-EYFP or SOM-Cre-Raptor-KO mice with a representative image of hChR2-mCherry expression, and diagram of behavioral testing sequence (open field and contextual fear conditioning). (E) Experimental protocol of TBS_opto_ given 30 min prior to contextual fear conditioning. (F) Left: Summary graph for all SOM-Cre-EYFP mice, showing reduced freezing in the long-term memory test in mice that received TBS_opto_ (n=8 mice, blue), relative to mice that received no TBS_opto_ (n=7 mice, violet) or TBS_opto_ without hChR2 (n=8 mice, gray). Mann-Whitney Rank Sum test (Pre vs Post conditioning), *** p < 0.001 (gray). Kruskal-Wallis One-way ANOVA on Ranks (Memory tests), Dunn’s pair-wise comparisons test, *** p=0.001, ns p>0.05. Right: Similar data presentation for SOM-Cre-Raptor-KO mice, showing significant freezing in the memory test relative to pre-conditioning, and no difference in freezing in the long-term memory test in mice that received TBS_opto_ (n=6 mice, blue) or no TBS_opto_ (n=7 mice, violet). Mann-Whitney Rank Sum test (Pre vs Post conditioning), ** p < 0.005 (gray). Unpaired t-test (Memory tests), ns p>0.05.

Next, we determined if LTP at PC-SOM synapses is sufficient to regulate hippocampus-dependent memory. We used CA1 injection of AAV2/9-CaMKIIa-hChR2(E123T/T159C)-mCherry in SOM-Cre-EYFP mice and the same TBS_opto_ induction protocol that elicit LTP at PC-SOM synapses in slices. We delivered the optogenetic stimulation in vivo and investigated its effect on contextual fear memory (Figure 3D and 3E). TBS_opto_ given in CA1 30 min prior to conditioning resulted in reduced freezing during the long-term contextual fear memory test, relative to mice receiving no TBS_opto_ prior to conditioning (73.7% reduction; Figure 3F) or TBS_opto_ in the absence of hChR2 (71.5% reduction; Figure 3F). Mice in the three groups showed similar normal anxiety level (time spent in periphery or center, and ratio of time in center/periphery; Figure S4E) and locomotion (total distance traveled and zone transitions; Figure S4F) in the open field test. These results indicate that the TBS_opto_ induction protocol that elicit LTP at PC-SOM synapses is sufficient to regulate long-term contextual fear memory in vivo.

Since LTP induced by TBS_opto_ at PC-SOM synapses, and its regulation of CA1 PC metaplasticity, are blocked by conditional knock-out of *Rptor* in SOM cells (Figures 1 and 2), we tested next if the reduction of fear memory by TBS_opto_ was due to mTORC1 signaling in SOM cells. TBS_opto_ in CA1 given 30 min prior to conditioning in SOM-Cre-Raptor-KO mice did not affect freezing during the long-term contextual fear memory test, relative to mice not receiving TBS_opto_ prior to conditioning (Figure 3F). Both mice groups showed increased freezing in the memory tests demonstrating significant contextual fear learning (Figure 3F) and behaved similarly in the open field test (Figure S4D-S4F) indicating normal anxiety level and locomotion. These results indicate that the TBS_opto_-induced impairment of fear memory was dependent on mTORC1 signaling in SOM interneurons and not due to other non-specific effects of optogenetic activation of CA1 PCs, suggesting that TBS_opto_-induced LTP at PC-SOM synapses is sufficient for the regulation of hippocampus-dependent memory.

### Prior induction of LTP by TBS_opto_ results in subsequent TBS- and learning-induced depotentiation

Since the behavior experiments indicate that TBS_opto_ impairs contextual fear memory consolidation when given prior to contextual fear conditioning, we examined the interaction between optogenetically- and electrically-induced LTP of electrically-evoked EPSPs in SOM cells in slices (Figure 4A).

**Figure 4.**
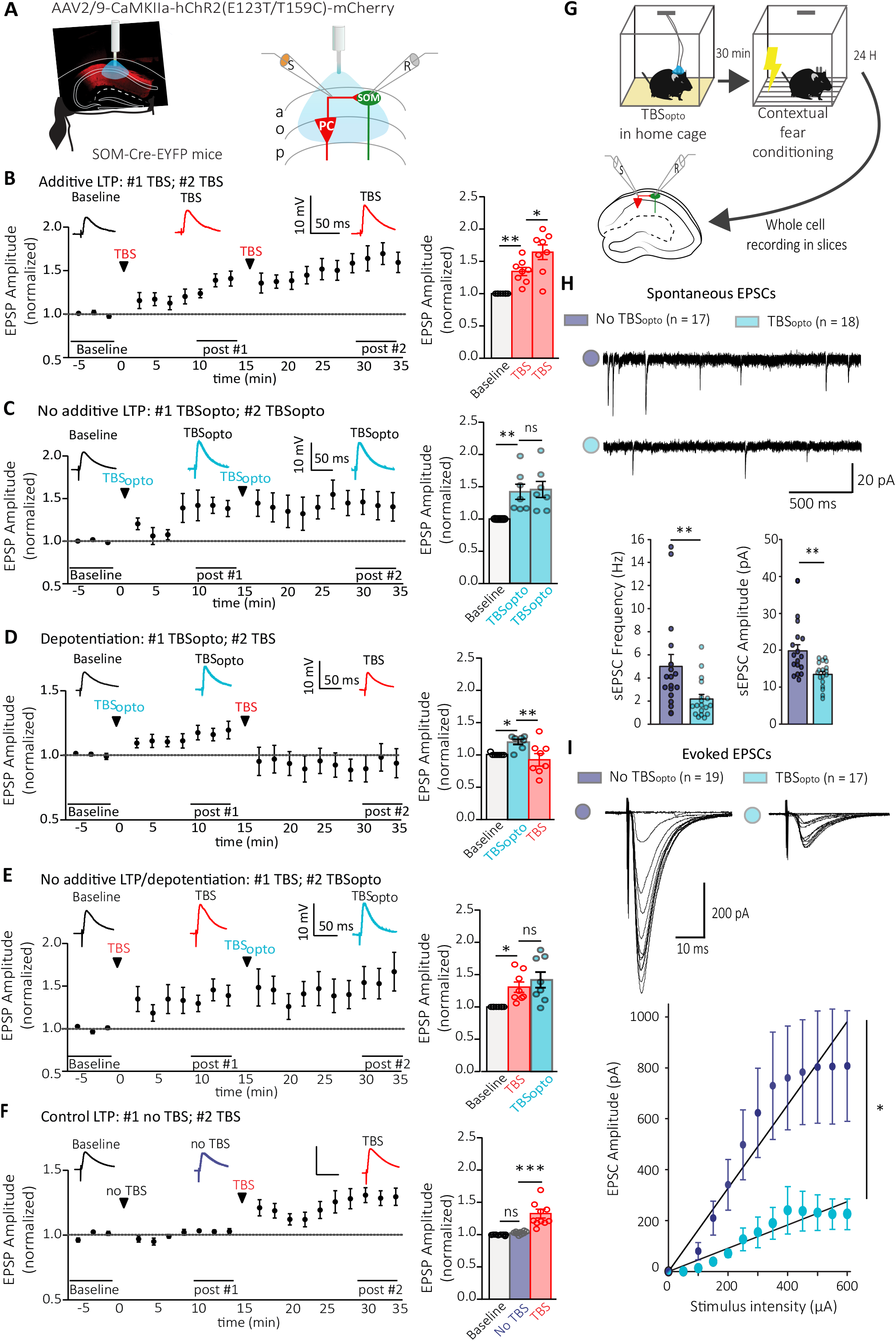
LTP induction by TBS_opto_ interferes with subsequent induction of Hebbian or learning-related LTP. (A) Left: schematic of experimental paradigm with viral injections in dorsal hippocampus of SOM-Cre-EYFP. Right: diagram of whole-field optogenetic stimulation of CA1 pyramidal cells (TBS_opto_), electrical stimulation of afferents in stratum oriens/alveus (S), and whole cell recording (R) of electrically-evoked EPSPs in SOM interneurons. (B-F) (Left) Time plots of electrically-evoked EPSP amplitude for all cells (with above, representative average EPSPs during baseline, post induction #1 and #2), and (right) summary bar graphs of EPSP amplitude for all cells during baseline, 10-15 min after induction #1 and 15-20 min after induction #2, showing: (B) LTP following a first episode of TBS of afferents (TBS#1) that is further increased after a second TBS episode (n = 8 cells); (C) LTP following a first episode of TBS_opto_ without further increase following a second TBS_opto_ episode (n = 7 cells); (D) LTP following a first episode of TBS_opto_ followed by depotentiation after a subsequent episode of TBS (n = 8 cells); (E) LTP following a first episode of TBS and absence of depotentiation after a subsequent TBS_opto_ episode (n = 8 cells); and (F) LTP following TBS given at the induction #2 time point only (n = 10 cells). rmANOVA, Tukey’s multiple comparisons test; * p < 0.05, ** p < 0.01, *** p < 0.0001, ns p > 0.05. (G) Diagram of experimental paradigm with in vivo optogenetic stimulation of CA1 pyramidal cells (TBS_opto_) prior to contextual fear conditioning, and ex vivo whole cell recording of spontaneous and electrically-evoked EPSCs (electrical stimulation in stratum oriens/alveus) in SOM interneurons in slices at 24 hours post conditioning. (H) Top: Spontaneous EPSCs from representative cells from mice receiving contextual fear conditioning only (violet) or TBS_opto_ prior to contextual fear conditioning (blue). Bottom: Summary bar graphs of sEPSC frequency and amplitude for all cells, showing decrease in frequency and amplitude of sEPSCs in cells from mice with TBS_opto_ prior to contextual fear conditioning (blue; n = 18 cells, 3 mice), relative to cells from mice with contextual fear conditioning only (violet; n = 17 cells, 4 mice). Mann-Whitney Rank Sum Test, ** p < 0.01. (I) Top: Family of EPSCs evoked by increasing intensity of stimulation in representative cells from mice receiving contextual fear conditioning only (violet) or TBS_opto_ prior to contextual fear conditioning (blue). Bottom: Summary plots of input-output function of EPSC amplitude as a function of stimulation intensity and linear fit (synaptic gain), showing that the input-output function of evoked EPSCs was decreased in cells from mice with TBS_opto_ prior to contextual fear conditioning (blue; n = 17 cells, 3 mice), relative to cells from mice that received contextual fear conditioning only (violet; n = 19 cells, 4 mice). Mann-Whitney Rank Sum Test, * p < 0.05.

After a baseline period, a first episode of electrical TBS elicited LTP of EPSP amplitude (134.0% of baseline at 10-15 min post-induction TBS#1), and a second TBS episode induced a further increase in LTP (164.0% of baseline at 30-35 min post-induction TBS#1; Figure 4B). In contrast, a first episode of TBS_opto_ elicited LTP of EPSP amplitude (140.0% of baseline at 10-15 min post-induction of TBS_opto_ #1), and a second TBS_opto_ episode did not induce a further increase in LTP (142.0% of baseline at 30-35 min post-induction TBS_opto_ #1; Figure 4C). Interestingly, after a first episode of TBS_opto_ that elicited LTP of EPSP amplitude (121.0% of baseline at 10-15 min post-induction of TBS_opto_ #1), a subsequent TBS episode induced depotentiation of EPSP amplitude (94.0% of baseline at 30-35 min post-induction TBS_opto_ #1; Figure 4D). Such a depotentiation was absent when a first episode of TBS (130.0% of baseline at 10-15 min post-induction of TBS #1) was followed by a subsequent TBS_opto_ episode (141.0% of baseline at 30-35 min post-induction TBS #1; Figure 4E). Finally, a single TBS episode given at the later time point induced LTP (129.0% of baseline; Figure 4F), suggesting that the depotentiation induced by TBS was not due to a response run-down due to recording time, or dialysis of cell content. Thus, prior induction of LTP by TBS_opto_ engages some mechanisms resulting in depotentiation when TBS is given subsequently.

Contextual fear conditioning induces a persistent LTP of PC-SOM synapses, that can be recorded ex vivo 24h after conditioning (Artinian et al., 2019). Therefore, we used ex vivo whole cell recordings in slices 24 hours after conditioning, to examine if in vivo TBS_opto_ given 30 min prior to contextual fear conditioning interferes with learning-induced LTP at excitatory synapses of SOM cells (Figure 4G). SOM cells from mice that received TBS_opto_ prior to contextual fear conditioning exhibited decreases in spontaneous EPSC frequency and amplitude, relative to cells from mice with contextual fear conditioning only (56.1% decrease in frequency, 31.9% decrease in amplitude; Figure 4H). Similarly, the input-output function of electrically-evoked EPSCs was decreased in SOM cells from mice that received TBS_opto_ prior to contextual fear conditioning, relative to cells from mice with contextual fear conditioning only (64.1% decrease in slope of EPSC input-output function; Figure 4I). These results suggest that in vivo TBS_opto_ given 30 min prior to contextual fear conditioning interferes with learning-induced LTP at PC-SOM synapses, consistent with our findings that induction of Hebbian LTP results in depotentiation at PC-SOM synapses after prior induction of LTP by TBS_opto_ in slices. Overall, our findings indicate that LTP at PC-SOM synapses is sufficient to regulate CA1 pyramidal cell metaplasticity and hippocampus-dependent memory.

## DISCUSSION

Long-term synaptic plasticity at excitatory input synapses onto specific inhibitory interneurons is an intriguing feature of hippocampal synaptic networks but its role in hippocampal memory function has remained largely undetermined (Kullmann et al., 2012, Honore et al., 2021, Bartos et al., 2011, McBain and Kauer, 2009). Brain-wide manipulation of mTORC1 activity specifically in SOM cells indicated that LTP at afferent synapses of hippocampal SOM cells contributes to regulation of CA1 network plasticity and long-term consolidation of spatial and contextual fear memory (Artinian et al., 2019). Here we used hippocampal CA1-specific optogenetic approaches to establish that LTP at PC-SOM synapses is sufficient to differentially control metaplasticity of SC-PC and TA-PC synapses, and regulate contextual fear memory, uncovering a long-term feedback mechanism controlling pyramidal cell synaptic plasticity and memory consolidation.

### TBS_opto_-induced LTP

TBS_opto_ was sufficient to induce LTP of synaptic responses elicited by optogenetic activation or local electrical stimulation of PC-SOM synapses. Optogenetically-induced LTP at PC-SOM synapses shared key features with Hebbian LTP induced by electrical tetanization: slow onset potentiation, frequency-dependence, mGluR1a-mediated, and SOM *versus* Pvalb cell-specificity (Perez et al., 2001, Lapointe et al., 2004, Croce et al., 2010, Vasuta et al., 2015). Since TBS_opto_ restricts activation to CA1 PCs, without affecting other CA1 afferent fibers potentially activated with tetanization by electrical stimulation (Nabavi et al., 2014, Nicholson and Kullmann, 2021, Cardin et al., 2010), selective activation of CA1 PC axons may, thus, be sufficient to induce Hebbian LTP at PC-SOM synapses. CA1 SOM cells receive other afferents, notably cholinergic projections from septum (Lovett-Barron et al., 2012, Leao et al., 2012), GABAergic fibers from septum (Sun et al., 2014), brainstem nucleus incertus (Szonyi et al., 2019) and local interneurons (Tyan et al., 2014), as well as noradrenergic, serotonergic and dopaminergic inputs (Pelkey et al., 2017, Oleskevich et al., 1989). Our findings establish that activation of these other projection systems is not required for LTP at PC-SOM synapses. This is in contrast to another form of plasticity at excitatory synapses onto putative-SOM cells in CA1 stratum oriens, anti-Hebbian LTP (Lamsa et al., 2007, Nicholson and Kullmann, 2021), which can be induced by optogenetic activation of cholinergic afferents but not of CA1 PCs, and, thus, is due to cholinergic heterosynaptic LTP at PC-oriens interneurons excitatory synapses (Nicholson and Kullmann, 2021). Our finding that TBS_opto_ of PC-SOM synapses is sufficient for induction of LTP is consistent with previous work that Hebbian LTP is afferent pathway-specific, occurring at CA1 PC-SOM synapses but not at CA3 PC-SOM synapses (Croce et al., 2010).

Interestingly, we uncovered that TBS_opto_-induced LTP at PC-SOM synapses was mTORC1-dependent and transcription-independent. A more persistent (lasting many hours) form of mGluR1a-mediated LTP at PC-SOM synapses is both transcription- and translation-dependent (Ran et al., 2009, Artinian et al., 2019). Since mTORC1 is a key signaling control mechanism of translation (Costa-Mattioli et al., 2009), our results suggest that mGluR1a-mediated LTP at PC-SOM synapse is translation-dependent. Thus, LTP at PC-SOM synapses may be mediated by mechanisms analogous to mGluR-mediated, mTORC1- and translation-dependent, transcription-independent LTD at SC-PC synapses (Huber et al., 2000, Banko et al., 2006). Such a mTORC1- and protein synthesis-dependent mechanism is consistent with the slow onset gradual development over minutes of LTP at PC-SOM synapses (Huber et al., 2000). SOM interneurons express both mGluR1 and mGluR5 (van Hooft et al., 2000, Topolnik et al., 2006). Previous work indicated that mGluR1a and mGluR5 signal via distinct pathways in SOM intermeurons, notably OLM cells (Topolnik et al., 2006). Activation of mGluR1a leading to Src/ERK-dependent Ca^2+^ entry via TRP channels and intracellular Ca^2+^ release are necessary for LTP induction at SOM interneuron excitatory synapses, whereas mGluR5 activation and intracellular Ca^2+^ release are not involved in LTP induction (Topolnik et al., 2006). Here we examined the role of mGluR1a in TBS_opto_-induced LTP at PC-SOM synapses, however additional work will be necessary to clarify if mGluR5 is implicated in this plasticity.

### Network regulation

Our finding that TBS_opto_ up-regulated LTP at SC-PC synapses and down-regulated LTP at TA- PC synapses in a mGluR1a- and SOM cell-mTORC1-dependent manner, is consistent with a differential gating of plasticity at PC input pathways by SOM cells (Leao et al., 2012) that is controlled over long-term periods by plasticity of SOM cell input synapses (Vasuta et al., 2015, Artinian et al., 2019, Sharma et al., 2020, Leao et al., 2012). It is noteworthy that TBS_opto_ did not affect basal transmission at SC- and TA-PC synapses, but regulated theta burst induced plasticity at these synapses. This modulation of plasticity reflects the property of PC-SOM synapses that show facilitation and postsynaptic firing with repeated stimulation, resulting in effective recruitment with repetitive activation of PCs (Croce et al., 2010, Honore et al., 2021, Pouille and Scanziani, 2004).

What are the network mechanisms responsible for the differential modulation of LTP of CA3 and entorhinal inputs of pyramidal cells? Previous work indicated that somatostatin-expressing oriens lacunosum-moleculare (OLM) interneurons differentially modulate CA3 and entorhinal inputs to CA1 pyramidal cells (Leao et al., 2012). OLM inhibitory interneurons suppress entorhinal inputs via postsynaptic inhibition of PC distal dendrites in stratum lacunosum-moleculare (Leao et al., 2012). In addition, OLM inhibitory interneurons facilitate CA3 inputs by disinhibition of the more proximal dendrites of pyramidal cells, i.e. OLM interneurons inhibit other inhibitory interneurons in stratum radiatum which themselves inhibit local pyramidal cell dendrites (Leao et al., 2012). Moreover, OLM interneurons similarly modulated LTP in these pathways, depressing LTP of entorhinal inputs and facilitating LTP of CA3 inputs (Leao et al., 2012). Our present findings are consistent with these mechanisms of regulation of CA1 network plasticity. In addition, previous work showed that a consequence of LTP at PC-SOM input synapses is to enhance synaptically-evoked firing of SOM interneurons (Croce et al., 2010). Thus, our present results and previous reports (Leao et al., 2012, Croce et al., 2010) are consistent with network mechanisms whereby LTP at input synapses of SOM INs increases their firing output, which enhances LTP of the CA3 input pathway via an increased disinhibitory action in proximal dendrites of pyramidal cells (Artinian et al., 2019, Vasuta et al., 2015), and suppresses LTP of the entorhinal input pathway via an increased inhibitory action in distal dendrites of pyramidal cells (Sharma et al., 2020).

Our findings highlight the different roles of long-term synaptic plasticity at excitatory synapses onto PC *versus* SOM interneurons. It has been proposed that LTP at PC excitatory synapses serves to generate enduring changes at synapses of engram cells that encode memories (Nabavi et al., 2014, Choi et al., 2018). In contrast, our results suggest that LTP at SOM cell excitatory synapses may serve to regulate the state of plasticity (or metaplasticity) of PC input synapses (Honore et al., 2021). These distinct roles suggest that hippocampal-dependent encoding of memories, via LTP in PCs, can still occur, albeit in a reduced manner, in the absence of LTP at SOM cell synapses, as for example in mice with conditional knockout of *Rptor* in SOM cells (Artinian et al., 2019). However, LTP at PC-SOM synapses controls the efficiency of hippocampal-dependent encoding memories by PCs, providing additional versatility to CA1 network plasticity. Our findings provide a link between LTP at PC-SOM synapses, regulation of metaplasticity at SC-PC and TA-PC synapses, and modulation of contextual fear memory. Contextual fear learning was previously shown to induce LTP at PC-SOM synapses (Artinian et al., 2019). Thus, during contextual fear learning, LTP is induced at PC-SOM synapses, and this may cause an upregulation of the CA1 network plasticity changes, such as LTP at SC-PC synapses, that encode memory. Thus, impairment in learning-induced plasticity at PC-SOM synapses caused by TBS_opto_ in vivo (Figure 4) may result in a loss of upregulation of metaplasticity in the CA1 network during learning (Figure 2), and in a deficit in memory consolidation (Figure 3). Such a link would be strengthened by a demonstration that contextual fear learning induces long-term changes at SC-PC and TA-PC synapses, and that these changes are modulated by in vivo manipulation of LTP at PC-SOM synapses. However, given the spatially restricted synaptic plasticity changes in pyramidal cells (Whitlock et al., 2006) and the sparse coding of synaptic changes in hippocampal engram cells (Choi et al., 2018), such experiments require different approaches that those used in the present study.

The gating of PC input pathways has been well established for OLM cells, a subtype of SOM cells (Leao et al., 2012). However, SOM cells also include other types of interneurons: bistratified cells, and other cells with both local projections and distal projections to septum, subiculum or retrohippocampal areas (CA3, dentate gyrus, entorhinal cortex) (Honore et al., 2021, Pelkey et al., 2017, Booker and Vida, 2018). Although excitatory synapses onto bistratified cells and OLM cells show Hebbian LTP (Perez et al., 2001, Croce et al., 2010), it remains to be determined if SOM projection cells also do. Thus, it will be interesting to build on our findings and determine how plasticity at input synapses of other CA1 SOM cell types may participate in network regulation, possibly even controlling more distant hippocampal-related pathways.

Learning-induced changes at the input synapses of SOM cells may only be part of a more integrated response of SOM cells during learning. An increase in the intrinsic excitability of CA1 SOM cells, due to a reduced Ca^2+^-dependent K^+^ conductance, was found after learning a hippocampus-dependent trace eyeblink conditioning task (McKay et al., 2013). Although it remains to be determined if SOM cell inhibitory synapses demonstrate learning-induced changes (Chiu et al., 2018, Udakis et al., 2020), mTORC1-dependent axonal sprouting by CA1 SOM cells takes place in the CACNA1A mouse model of epilepsy which is characterized by impaired synaptic inhibition by Pvalb interneurons (Jiang et al., 2018). These previous works and our findings indicate that multiple plasticity mechanisms occur in SOM cells at the level of their input function, intrinsic excitability and perhaps even output synapses, suggesting that an integrated gain-of-function response may occur in SOM cells during learning.

### TBS_opto_ interaction with Hebbian and learning-induced LTP

Our results that TBS_opto_ given *in vivo* prior to contextual fear conditioning affects long-term contextual fear memory suggests that LTP at PC-SOM synapses regulates hippocampal memory. In these experiments, TBS_opto_ was given in vivo. Thus, it is conceivable that optogenetic activation of PCs in the CA1 region also activated the other major synaptic target of pyramidal cells, subicular pyramidal neurons (Cenquizca and Swanson, 2007, Taube, 1993), inducing long-term plasticity at these synapses (O’Mara et al., 2000, Huang and Kandel, 2005) and influencing hippocampal memory function. To examine this possibility, we examined the effects of TBS_opto_ in mice with a cell-specific conditional knockout of *Rptor* in somatostatin interneurons, in which LTP at PC-SOM synapses is blocked. We found that the regulation of long-term contextual fear memory by TBS_opto_ was absent in these mice (Figure 3), suggesting that the regulation of hippocampal memory by TBS_opto_ was due to changes at PC-SOM synapses and not at other efferent targets of CA1 PCs, like subicular neurons. However, it would be interesting in future experiments to design an induction protocol applicable *in vivo* that selectively activates CA1 PC-SOM synaptic plasticity and, thus, rule out actions via other synaptic targets of CA1 PCs. Nonetheless, our results are consistent with our other observations that PC-SOM synapse efficacy was reduced, like fear memory, by TBS_opto_ given prior to learning (Figure 4), as well as with our previous findings that learning-induced LTP at PC-SOM synapses is necessary for long-term contextual fear memory (Artinian et al., 2019). It is noteworthy that TBS_opto_ in slices produces LTP at PC-SOM synapses and not at PC-PV synapses, indicating some CA1 PC target cell-specificity in TBS_opto_ induced long-term plasticity.

Our results indicate that LTP at PC-SOM synapses induced by TBS_opto_ share similar mechanisms as Hebbian LTP induced by electrical TBS and LTP induced by contextual fear learning at excitatory synapses onto SOM cells. As reported for LTP at hippocampal synapses (Bliss and Lomo, 1973), LTP induced by a first episode of TBS_opto_ occluded further LTP by a second TBS_opto_. However, LTP induced by a first episode of electrical TBS did not, and more LTP was induced by a second episode of TBS. In contrast, when electrical TBS was given after LTP induced by a first TBS_opto_ episode, depotentiation occurred at PC-SOM synapses. Similarly, in vivo, when contextual fear conditioning was given 30 min after TBS_opto_, the efficacy of PC-SOM synapses was reduced during *ex vivo* recordings 24 hours later. These results indicate that although they share similar mechanisms, LTP induced by TBS_opto_, electrical TBS and fear learning are not identical. This is not surprising given that electrical stimulation may activate additional intra-hippocampal fibers, and learning may implicate other systems in addition to CA1 PC-SOM synapses. These additional mechanisms are likely implicated in the depotentiation of PC-SOM synapses. Multiple mechanisms regulate PC-SOM synapse function and plasticity, notably pre- and post-synaptic GABA_B_-receptor mediated inhibition (Booker et al., 2020, Booker et al., 2018), astrocyte-mediated regulation of glutamate uptake (Huang et al., 2004), and GABA_A_ synaptic inhibition by vasoactive intestinal polypeptide inhibitory interneurons (Tyan et al., 2014). It will be important to determine if any of these control mechanisms are implicated in the modulation/depotentiation of PC-SOM synapses. Importantly, our findings that learning-induced potentiation is impaired by prior application of TBS_opto_ *in vivo* may explain why contextual fear memory may be impaired by TBS_opto_, providing further support that LTP induced at PC-SOM synapses is sufficient to regulate CA1 network metaplasticity and the contextual fear memory.

### Limitations of the study

Some limitations should be considered when interpreting our results. First, although our interpretation of the results is concordant with optical stimulation resulting in LTP at PC-SOM synapses *in vivo*, this remains to be demonstrated directly. This would require recording synaptic activity of SOM INs *in vivo* and assessing the effect of optogenetic induction *in vivo* on this synaptic activity. Such experiments however require different experimental approaches than those used here. Instead, we addressed this issue using the approach shown in Figure 4, with optical stimulation *in vivo*, contextual fear conditioning and *ex vivo* recordings. Our *ex vivo* results are consistent with optical stimulation *in vivo* eliciting LTP and with subsequent contextual fear conditioning resulting in depotentiation, instead of the late form of LTP that is normally elicited (Artinian et al., 2019). These results are analogous to the effect found in the slice experiments whereby optical stimulation *in vitro* elicits LTP and subsequent TBS with electrical stimulation results in depotentiation, instead of the LTP normally elicited. Thus, our results are consistent with the notion that optical stimulation elicits LTP at PC-SOM synapses both *in vivo* and *in vitro*. Previous work showed that a different induction protocol with repeated episodes of stimulation elicits a late, persistent form of LTP that lasts many hours to a day at SOM interneuron excitatory synapses (Artinian et al., 2019, Ran et al., 2009). Thus, it would be important to test if such a repeated optical stimulation protocol *in vivo* can produce persistent changes at PC-SOM synapses that can be detected with *ex vivo* recordings.

Second, a caveat of our study concerns our results that the regulation of long-term contextual fear memory by TBS_opto_ was absent in mice with conditional knockout of *Rptor* in somatostatin cells, suggesting that the regulation of hippocampal memory by TBS_opto_ was due to changes at PC-SOM synapses. The lack of effect of TBS_opto_ on contextual fear memory in the Raptor-KO model could be due to the low level of freezing response in these mice, indicating that these animals cannot learn any worse. In our experiments we used a weak contextual fear conditioning protocol based on previous work showing that inactivating SOM interneurons impairs contextual fear memory (Lovett-Barron et al., 2014). It is important to note that with this contextual fear conditioning protocol, SOM-Cre-Raptor-KO mice show impaired fear memory relative to control mice, however the SOM-Cre-Raptor-KO mice do display significant freezing responses in the memory probe test relative to pre-conditioning, indicating some level of contextual memory (Figure 3F). Using a stronger contextual fear conditioning protocol, previous work showed also that although the SOM-Cre-Raptor-KO mice display a memory impairment, they still show significant contextual fear learning (Artinian et al., 2019). Thus, these findings suggest that a component of contextual fear learning requires mTORC1 and SOM interneurons, but another component does not. A similar conclusion was found for spatial learning in the Barnes maze (Artinian et al., 2019). Thus, repeating the present experiments with a stronger contextual fear conditioning protocol (Artinian et al., 2019) could help resolve the possibility of a floor effect in our experiments. However, the residual significant freezing responses in SOM-Cre-Raptor-KO mice in the contextual fear memory probe test suggest the absence of a floor effect, and, thus, are consistent with the effects of TBS_opto_ on contextual fear memory being due to LTP at PC-SOM synapses.

## Supporting information

Star methods

## ACKNOWLEDGMENTS

This work was supported by grants to J.C.L. from the Canadian Institutes of Health Research (CIHR MOP-125985 and PJT-153311) and a Research Centre grant (Centre Interdisciplinaire de Recherche sur le Cerveau et l’Apprentissage; CIRCA) from the Fonds de la Recherche du Québec – Santé (FRQS). J.C.L. is the recipient of the Canada Research Chair in Cellular and Molecular Neurophysiology (CRC 950-231066) and a member of the Research Group on Neural Signaling and Circuitry (GRSNC) at Université de Montréal.

## AUTHOR CONTRIBUTIONS

Conceptualization, J.C.L., A.A. and E.H.; Methodology, A.A., E.H and I.L.; Software, E.H.; Formal Analysis, A.A. and E.H.; Investigation, A.A., E.H., I.L. and F.X.M.; Writing – Original Draft, J.C.L., A.A., E.H. and I.L.; Writing – Review & Editing, J.C.L., A.A. and E.H.; Visualization, A.A., E.H. and I.L.; Supervision, Project Administration, and Funding Acquisition, J.C.L.

## STAR METHODS

### RESOURCE AVAILABILITY

#### Lead Contact

Further information and requests for resources should be directed to and will be fulfilled by the Lead Contact, Dr. Jean-Claude Lacaille.

#### Materials Availability

This study did not generate new unique reagents.

#### Data and Code Availability

The published article includes all datasets generated or analyzed during this study. The raw electrophysiology data supporting the current study have not been deposited in a public repository because there is currently no standardized format or repository for such data, but they are available from the corresponding author on request.

### EXPERIMENTAL MODEL AND SUBJECT DETAILS

#### Animals

Experimental protocols were approved by the Animal Care Committee at the Université de Montréal (Comité de Déontologie de l’Expérimentation sur les Animaux; CDEA Protocols # 17-001, 17-002, 18-002, 18-003, 19-003, 19-004, 20-001, 20-002, 21-001, 21-002) and experiments were performed in accordance with the Canadian Council of Animal Care guidelines.

*Sst*^ires-Cre^ mice (Taniguchi et al., 2011) were crossed with *Rosa26*^lsl-EYFP^ reporter mice (Madisen et al., 2010) to generate Cre-dependent Enhanced Yellow Fluorescent Protein (EYFP) expression in SOM interneurons (SOM-Cre-EYFP mice) (Artinian et al., 2019). *Sst*^ires-Cre^;*Rosa26*^lsl-EYFP^ mice were crossed with floxed *Rptor* mice (Sengupta et al., 2010) for cell-specific homozygous knock-out of *Rptor* in SOM cells (SOM-Cre-Raptor-KO mice) (Artinian et al., 2019). *Pvalb^ires-^*^Cre^ mice (Hippenmeyer et al., 2005) were used for Cre-dependent targeting of PV interneurons (Pvalb-Cre mice). Mice were housed 2-4 animals per cage, except for *in vivo* optogenetic studies in which mice were housed singly after cannula implantation. Food and water were *given ad libitum*. The mice were maintained on a 12 h light/dark cycle with all experiments performed during the light phase.

The *in vitro* electrophysiological slice experiments were performed on 6-7 weeks old male or female mice at the diestrus phase to reduce possible variability in responses. Behavioral studies and *ex vivo* electrophysiological slice experiments were carried out on 8-10 weeks old male mice.

### METHOD DETAILS

#### Virus injection

Four weeks old mice were given an intraperitoneal (IP) injection of ketamine (50 mg/kg i.p.) and xylazine (5 mg/kg i.p.) and placed in a stereotaxic frame (Stoelting). For *in vitro* slice experiments, AAV2/9-CaMKIIa-hChR2(E123T/T159C)-mCherry (1.5-1.77×10^12^ particles/ml) or AAV2/9-CaMKIIa-mCherry (2.20×10^13^ particles/ml) were injected bilaterally in CA1 hippocampus (coordinates relative to bregma: −1.95 mm AP, ±1.3 mm ML, and −1.3 mm DV) of mice of either sex. Viral solution (0.5-0.8μl) was delivered at a flow rate of 50-100 nl/min using a 10μl Hamilton syringe and a microfluidic pump (Harvard Apparatus). The needle was left in place for at least 5 minutes after injection. For some slice experiments, Pvalb-Cre mice were injected, in addition, with AAV2/9-EF1a-DIO-EYFP (3.95×10^13^ particles/ml). For *in vivo* optogenetic experiments, 7-8 weeks old male mice were used and treated as above. In some *in vivo* experiments, mice were injected bilaterally 0.5μl of AAV2/9-flex-Arch-GFP (1.74-1.83×10^13^ particles/ml) or AAV2/9-EF1a-DIO-EYFP (3.95×10^13^ particles/ml) as above.

#### Hippocampal slice preparation

Six to seven weeks old animals (10-18 days after viral injection) were anesthetized by isoflurane inhalation and the brain was rapidly removed and placed in cold (4°C) sucrose-based cutting solution saturated with 95% O_2_ and 5% CO_2_ containing the following (in mM): 87 NaCl, 2.5 KCl, 1.25 NaH_2_PO_4_, 7 MgSO_4_, 0.5 CaCl_2_, 25 NaHCO_3_, 25 glucose, 9.5 ascorbic acid, 3.0 pyruvic acid and 75 sucrose, pH 7.4, and 295-300 mOsmol/L. A block of tissue containing the hippocampus was prepared and transverse hippocampal slices (300 and 400 μm thickness for whole-cell and field recordings, respectively) were cut with a vibratome (Leica VT1000S). Slices were transferred to artificial CSF (ACSF; saturated with 95% O_2_ and 5% CO_2_) containing the following (in mM): 124 NaCl, 2.5 KCl, 1.25 NaH_2_PO_4_, 4 MgSO_4_, 4 CaCl_2_, 26 NaHCO_3_, and 10 glucose (pH 7.3–7.4, and 295–300 mOsmol/L) at 32°C for 30 minutes, and afterward maintained at room temperature (20–22°C) for at least 90 minutes and until recording experiments. Individual slices were transferred to a submersion chamber perfused (3-4 ml/min) with ACSF at 31 ± 0.5°C, and with CA1 and CA3 regions separated by a surgical cut. Synaptic inhibition was intact in most experiments, except for voltage clamp recordings of EPSCs (see below). For *ex vivo* recordings after learning, slices were obtained as above from animals 24 hours after optogenetic stimulation and contextual fear conditioning.

#### Whole-cell patch clamp recording

EYFP-expressing CA1 interneurons were identified using an upright microscope (Nikon Eclipse, E600FN), equipped with a water-immersion long-working distance objective (40x; differential interference contrast, DIC), epifluorescence and an infrared CCD camera (CXE-B013-U; Mightex). Whole-cell current-clamp recordings were obtained using borosilicate glass pipettes (3-5 MΩ) filled with intracellular solution containing (in mM): 120 KMeSO_4_, 10 KCl, 0.5 EGTA, 10 HEPES, 2.5 MgATP, 0.3 NaGTP and 10 Na_2_-phosphocreatine (pH 7.3, 290-298 mOsmol/L). For whole-cell voltage-clamp recordings, the intracellular solution contained (in mM): 120 CsMeSO_3_, 5 CsCl, 2 MgCl_2_, 10 HEPES, 0.5 EGTA, 10 Na_2_-phosphocreatine, 2 ATP-Tris, 0.4 GTP-Tris, 0.1 spermine, 2 QX314 (pH 7.2–7.3, 300 mOsmol/L). Data was acquired using a Multiclamp 700B amplifier (Molecular Devices) and digitized at 10-20 kHz using Digidata 1440A and pClamp 10.5/10.7 software (Molecular Devices). Recordings were low-pass filtered at 2 kHz and membrane potential was corrected for liquid junction potential (11 mV). Access resistance was monitored throughout experiments and data were included only if the holding current and series resistance were stable (changes <20% of initial value).

#### Whole cell recording of synaptic responses

EPSPs and EPSCs evoked by electrical stimulation were elicited using constant current pulses (50 μs duration) via an ACSF-filled bipolar theta-glass electrode positioned approximately 100-150 μm lateral from the recorded cell soma in the alveus near the border with CA1 stratum oriens. EPSCs were recorded in the presence of DL-2-Amino-5-phosphonopentanoic acid (DL-AP5; 50 μM) and SR-95531 (Gabazine, 5 μM) to block NMDA and GABA_A_ receptors, respectively. Input-output function of EPSCs was studied by delivering current pulses of incremental intensity (0–600μA, 50μA steps); 10 trials per pulse intensity were delivered and responses averaged to determine EPSC amplitude (initial EPSC peak). The slope (synaptic gain) and *x*-intercept (minimal stimulation intensity) of the linear regression of the input-output relationship were measured on averaged responses of individual cells. Spontaneous EPSCs were recorded over a period of 5 min and generally 300 consecutive events were analyzed (except for some cells with low frequency of events: TBS_opto_ group 4 cells with 168, 188, 200 and 261 events; no TBS_opto_ group 2 cells with 279 and 294 event) for frequency and amplitude (Clampfit 10.7).

EPSPs evoked by focal optogenetic stimulation through the objective were elicited using a Polygon400 Multiwavelength Dynamic Patterned Illuminator (Mightex). An area (20×20 to 20×30 μm) lateral to the recorded cell soma was illuminated with brief pulses of blue light (470 nm; duration 0.5-2 ms). EPSPs evoked by whole field optogenetic stimulation were elicited using a 4-Channel LED Driver (DC4104; Thorlabs) coupled to an optical fiber (MF79L01; 400 μm core diameter, 0.39 NA; Thorlabs) and fiber optic cannula (CFM14L20; 400 μm core diameter, 0.39 NA; Thorlabs) positioned above the CA1 stratum oriens region of the slice with brief light pulses (470 nm; duration 0.5-2 ms). In some experiments, the mGluR1a antagonist LY367385 (40 μM) was added in the ACSF during recordings. In other experiments, slices were pre-incubated in ACSF with 2 μM of the irreversible transcription inhibitor actinomycin D for 15 min prior to recording (Yuan and Burrell, 2013, Younts et al., 2016).

#### LTP induction protocol during whole cell current clamp recording

Test EPSPs were evoked at 0.1Hz. LTP induced by electrical stimulation was elicited by theta burst stimulation (TBS) consisting of 3 episodes (given at 30 s intervals) of 5 bursts (at 250 ms interburst intervals) of 4 electrical pulses at 100 Hz (Vasuta et al., 2015). LTP induced by optogenetic stimulation of CA1 pyramidal cells (TBS_opto_) was elicited by TBS consisting of 3 episodes (given at 30 s intervals) of 5 bursts (at 300 ms inter-burst interval) of 4 light pulses at 80 Hz. In some experiments, low frequency optogenetic stimulation (LFS_opto_) was given as 3 episodes (given at 30 s intervals) of 3 bursts (at 500 ms inter-burst interval) of 3 light pulses at 20 Hz. Test EPSP amplitude was averaged in 2 minutes bins before and after LTP induction. EPSPs were characterized in one cell per slice, and responses were analyzed off-line using Clampfit (pClamp 10.5/10.6; Molecular Devices) and GraphPad Prism 6.

#### Field potential recording

For experiments with field potential recordings, transverse hippocampal slices were prepared as described above, except oxygenated ACSF contained 1.3 mM MgSO_4_ and 2.5 mM CaCl_2_. The slices were let to recover for at least 2 h at 32°C in ACSF, and for an additional 30 minutes at 27°–28°C while submerged in a recording chamber continuously perfused (2–2.5 ml/min) with ACSF. Extracellular field EPSPs were recorded with borosilicate glass pipettes (3–4MΩ) filled with ACSF using a differential extracellular amplifier (Microelectrode AC Amplifier Model 1800, A-M Systems), filtered at 2 kHz, digitized at 10 kHz (Digidata 1440A), and analyzed with pClamp10.5. For Schaffer collateral-evoked fEPSPs, the concentric bipolar tungsten stimulating electrode (FHC) and recording pipette were placed in stratum radiatum. For temporo-ammonic pathway evoked fEPSPs, the stimulation and recording electrodes were positioned in stratum lacunosum-moleculare. Stimulus (0.1 ms duration; 15 sec^−1^) strength was adjusted to elicit 50% of maximal fEPSP and fEPSP slope was measured at 10–90% of fEPSP amplitude. CA1 pyramidal cell long-term potentiation was induced by weak theta burst stimulation (wTBS; 2 bursts of 4 pulses at 100 Hz, with 200 ms inter-burst interval) (Leao et al., 2012). Optogenetic induction of LTP in SOM interneurons was elicited as described above through a fiber optic positioned above the CA1 stratum oriens/alveus region of the slice.

#### Optogenetic stimulation in vivo

One week after viral injections, mice were anesthetised using ketamine and xylazine as described above, and fiber cannulas were stereotaxically implanted bilaterally (coordinates relative to bregma: −1.95 AP; ±1.60 ML; and −1.15 DV) and sealed with dental cement (Metabond, Parkell Inc). Animals were housed singly after the optic fiber implantation and allowed to rest for a week before behavioral experimentation. For in vivo optogenetic stimulation, a Quadruple Laser Diode Fiber Light Source (LDFLS_405/100_450/070_520/060_638/080; Doric Lenses Inc) was coupled to Mono Fiberoptic Patchcords (MFP_200/240/900-0.22_2m_FC-ZF1.25(F), 200 μm Core diameter, 0.22 NA; Doric Lenses Inc) and hand-made fiber optic cannulas (optic fiber: FT200EMT, 200 μm Core diameter, 0.22 NA; ceramic ferrule: CFLC230-10; Thorlabs).

#### Behavioral experiments

For all behavioral experiments, the experimenter was blind to the experimental groups (opsin *versus* reporter) from the day habituation began until data analysis was completed.

##### Habituation

Mice were gently handled daily for 5 days (5min/day) to progressively habituate them to the experimenter and reduce the stress related to the experimental handling and optic fiber connection (without light stimulation).

##### Open field test

Mice were allowed to freely explore a square arena (45 x 45 cm; PanLab) for 15 min for the Arch experiments and 5 min for the hChR2 experiments. The anxiety level was assessed by measuring the time spent and the distance travelled in the center (1/3 central zone) of the arena compared to the periphery (1/3 peripheral zone). Locomotion was evaluated by measuring the total distance traveled and the number of zone transitions (16 equal squared zones). Mice were first video-tracked at 25 frames/s and their movements subsequently analyzed using a position tracking system (Smart 3.0; PanLab). For the Arch experiments, mice received continuous light stimulation (520 nm) for the 5-10 min period.

#### Contextual fear conditioning

Mice were trained in conditioning chambers that were housed in sound- and light-isolated cubicles (Coulbourn Instruments). The chambers contained a stainless-steel grid floor, overhead LED lighting, camera and were supplied with background noise (60 dB) by an air extractor fan. The experimental protocol was based on Lovett-Baron and coworkers (Lovett-Barron et al., 2014) with slight modifications. The training context was rectangular with 2 transparent and 2 stainless steel walls, was cleaned with 70% ethanol solution before and after each trial, and ethanol solution was added under the grid floor as a contextual smell. After 2 min free exploration, the animals received a weak fear conditioning consisting of 2 presentations of unconditioned stimulus (2 sec foot shock, 0.8mA; separated by 60 sec). To test for long-term contextual fear memory, the mice were returned to the training context during a test period of 3 min, at 24 h after conditioning. The freezing behavior was assessed using FreezeFrame (Coulbourn Instruments).

For the Arch optogenetic experiments, mice received light stimulation (520 nm; 6 sec pulses) in phase (starting 2 sec before) or shifted (30 sec before) relative to foot shocks during conditioning. For the TBS optogenetic experiments, mice were brought in the testing room on the training day and, in their home cage, given the optogenetic TBS protocol (473 nm; 5 bursts of 4 light pulses of 1ms duration at 80 Hz, given at 300 ms interburst interval, and repeated 3 times with 30 s interval) or no light stimulation, 30 min prior to contextual fear conditioning.

#### Histology

Following *in vivo* optogenetic experiments, mice were deeply anesthetized intraperitoneally with sodium pentobarbital (MTC Pharmaceuticals), perfused transcardially first with 0.1M phosphate buffer saline (PBS) and next with 4% paraformaldehyde in 0.1M PBS (PFA). Brains were isolated, postfixed in 4% PFA for 24 hours and cryoprotected in 30% sucrose. Coronal brain sections (50 μm thick) were obtained using freezing microtome (SM200R; Leica) and mounted in ProLong Gold (Invitrogen). Sections were examined using an epifluorescence microscope (Eclipse E600; Nikon) and images were acquired with the Simple PCI software.

Immunofluorescence was used to enhance GFP fluorescence associated with Arch. For these mice, brain sections were obtained as described above, permeabilized with 0.5% Triton X-100 in PBS for 15 minutes, and unspecific binding was blocked with 10% normal goat serum in 0.1% Triton X-100/PBS for 1 hour. Rabbit polyclonal GFP (1/200) antibodies were incubated for 48 hours at 4°C. Sections were subsequently incubated at room temperature with Alexa Fluor 594-conjugated goat anti-rabbit IgG (1/500; 90 min), mounted in ProLong Gold, and imaged as described above. For behavioral experiments, data were excluded if mice did not show virus expression restricted to CA1, and if optic fibers placement was outside the CA1 hippocampus.

### QUANTIFICATION AND STATISTICAL ANALYSIS

No statistical methods were used to predetermine the sample size, but our sample sizes are similar to those used in the field. Statistical analysis was performed using SigmaPlot or GraphPad Prism 6 and 9. Data were tested for normality using the Shapiro–Wilk, D’Agostino and Pearson, Kolmogorov-Smirnov, and Brown–Forsythe tests. We used Student’s *t* tests, one- or two-way ANOVA, and one- or two-way mixed repeated-measures ANOVA, with Tukey’s pairwise comparison tests (with Bonferroni adjustments for multiple comparisons) or with Holm-Sidak pairwise comparison tests, when data passed normality and homoscedasticity assumptions. Wilcoxon matched pairs signed rank tests, Mann–Whitney Rank Sum tests and Kruskal-Wallis one-way ANOVA on Ranks with Dunn’s pairwise comparisons, were used when data did not pass normality and homoscedasticity assumptions. All the tests were two-sided. All data in the Figures are presented as mean ± SEM. Asterisks in Figures denote statistical significance levels for specified tests (* *p* < 0.05; ** *p* < 0.01; *** *p* < 0.001; ns, not significant). Detailed results of all statistical tests referenced per Figure panel are given in Supplemental Table 1. Baseline EPSP amplitudes are reported for all groups in Supplemental Table 2, with Dunn’s multiple comparisons tests indicating no differences between SOM interneuron groups.

## Supplemental Information

**Table S1. Details of statistical comparisons (Excel file), related to Figures 1–4, S1-4.**

**Table S2. Light-evoked EPSP and electrical-evoked EPSP amplitude pre-TBS in all related experiments, related to Figures 1, 4, S2 and S3.**

**Figure S1. Optogenetically-induced LTP at PC-SOM synapses in representative cells, related to Figure 1.**

**Figure S2. Optogenetically-induced LTP of electrically-evoked EPSPs in SOM cells, related to Figure 1.**

**Figure S3. Whole-field TBS_opto_-induced LTP at PC-SOM synapses and differential regulation of LTP at SC-PC and TA-PC synapses, related to Figure 2.**

**Figure S4. Normal anxiety and locomotion in open field tests, related to Figure 3.**

## REFERENCES

Artinian, J., Jordan, A., Khlaifia, A., Honore, E., La Fontaine, A., Racine, A. S., Laplante, I. & Lacaille, J. C. 2019. Regulation of Hippocampal Memory by mTORC1 in Somatostatin Interneurons. J Neurosci, 39, 8439–8456.

Banko, J. L., Hou, L., Poulin, F., Sonenberg, N. & Klann, E. 2006. Regulation of eukaryotic initiation factor 4E by converging signaling pathways during metabotropic glutamate receptor-dependent long-term depression. J Neurosci, 26, 2167–73.

Bartos, M., Alle, H. & Vida, I. 2011. Role of microcircuit structure and input integration in hippocampal interneuron recruitment and plasticity. Neuropharmacology, 60, 730–9.

Bezaire, M. J. & Soltesz, I. 2013. Quantitative assessment of CA1 local circuits: knowledge base for interneuron-pyramidal cell connectivity. Hippocampus, 23, 751–85.

Bliss, T. V. & Lomo, T. 1973. Long-lasting potentiation of synaptic transmission in the dentate area of the anaesthetized rabbit following stimulation of the perforant path. J Physiol, 232, 331–56.

Booker, S. A., Harada, H., Elgueta, C., Bank, J., Bartos, M., Kulik, A. & Vida, I. 2020. Presynaptic GABAB receptors functionally uncouple somatostatin interneurons from the active hippocampal network. Elife, 9.

Booker, S. A., Loreth, D., Gee, A. L., Watanabe, M., Kind, P. C., Wyllie, D. J. A., Kulik, A. & Vida, I. 2018. Postsynaptic GABABRs Inhibit L-Type Calcium Channels and Abolish Long-Term Potentiation in Hippocampal Somatostatin Interneurons. Cell Rep, 22, 36–43.

Booker, S. A. & Vida, I. 2018. Morphological diversity and connectivity of hippocampal interneurons. Cell Tissue Res, 373, 619–641.

Cardin, J. A., Carlen, M., Meletis, K., Knoblich, U., Zhang, F., Deisseroth, K., Tsai, L. H. & Moore, C. I. 2010. Targeted optogenetic stimulation and recording of neurons in vivo using cell-type-specific expression of Channelrhodopsin-2. Nat Protoc, 5, 247–54.

Cenquizca, L. A. & Swanson, L. W. 2007. Spatial organization of direct hippocampal field CA1 axonal projections to the rest of the cerebral cortex. Brain Res Rev, 56, 1–26.

Chiu, C. Q., Martenson, J. S., Yamazaki, M., Natsume, R., Sakimura, K., Tomita, S., Tavalin, S. J. & Higley, M. J. 2018. Input-Specific NMDAR-Dependent Potentiation of Dendritic GABAergic Inhibition. Neuron, 97, 368–377 e3.

Choi, J. H., Sim, S. E., Kim, J. I., Choi, D. I., Oh, J., Ye, S., Lee, J., Kim, T., Ko, H. G., Lim, C. S. & Kaang, B. K. 2018. Interregional synaptic maps among engram cells underlie memory formation. Science, 360, 430–435.

Costa-Mattioli, M., Sossin, W. S., Klann, E. & Sonenberg, N. 2009. Translational control of long-lasting synaptic plasticity and memory. Neuron, 61, 10–26.

Croce, A., Pelletier, J. G., Tartas, M. & Lacaille, J. C. 2010. Afferent-specific properties of interneuron synapses underlie selective long-term regulation of feedback inhibitory circuits in CA1 hippocampus. J Physiol, 588, 2091–107.

Freund, T. F. & Buzsaki, G. 1996. Interneurons of the hippocampus. Hippocampus, 6, 347–470.

Fuhrmann, F., Justus, D., Sosulina, L., Kaneko, H., Beutel, T., Friedrichs, D., Schoch, S., Schwarz, M. K., Fuhrmann, M. & Remy, S. 2015. Locomotion, Theta Oscillations, and the Speed-Correlated Firing of Hippocampal Neurons Are Controlled by a Medial Septal Glutamatergic Circuit. Neuron, 86, 1253–64.

Hippenmeyer, S., Vrieseling, E., Sigrist, M., Portmann, T., Laengle, C., Ladle, D. R. & Arber, S. 2005. A developmental switch in the response of DRG neurons to ETS transcription factor signaling. PLoS Biol, 3, e159.

Honore, E., Khlaifia, A., Bosson, A. & Lacaille, J. C. 2021. Hippocampal Somatostatin Interneurons, Long-Term Synaptic Plasticity and Memory. Front Neural Circuits, 15, 687558.

Huang, Y. H., Sinha, S. R., Tanaka, K., Rothstein, J. D. & Bergles, D. E. 2004. Astrocyte glutamate transporters regulate metabotropic glutamate receptor-mediated excitation of hippocampal interneurons. J Neurosci, 24, 4551–9.

Huang, Y. Y. & Kandel, E. R. 2005. Theta frequency stimulation up-regulates the synaptic strength of the pathway from CA1 to subiculum region of hippocampus. Proc Natl Acad Sci U S A, 102, 232–7.

Huber, K. M., Kayser, M. S. & Bear, M. F. 2000. Role for rapid dendritic protein synthesis in hippocampal mGluR-dependent long-term depression. Science, 288, 1254–7.

Jiang, X., Lupien-Meilleur, A., Tazerart, S., Lachance, M., Samarova, E., Araya, R., Lacaille, J. C. & Rossignol, E. 2018. Remodeled cortical inhibition prevents motor seizures in generalized epilepsy. Ann Neurol, 84, 436–451.

Katona, L., Lapray, D., Viney, T. J., Oulhaj, A., Borhegyi, Z., Micklem, B. R., Klausberger, T. & Somogyi, P. 2014. Sleep and movement differentiates actions of two types of somatostatin-expressing GABAergic interneuron in rat hippocampus. Neuron, 82, 872–86.

Klausberger, T., Magill, P. J., Marton, L. F., Roberts, J. D., Cobden, P. M., Buzsaki, G. & Somogyi, P. 2003. Brain-state- and cell-type-specific firing of hippocampal interneurons in vivo. Nature, 421, 844–8.

Klausberger, T. & Somogyi, P. 2008. Neuronal diversity and temporal dynamics: the unity of hippocampal circuit operations. Science, 321, 53–7.

Kullmann, D. M., Moreau, A. W., Bakiri, Y. & Nicholson, E. 2012. Plasticity of inhibition. Neuron, 75, 951–62.

Lamsa, K. P., Heeroma, J. H., Somogyi, P., Rusakov, D. A. & Kullmann, D. M. 2007. Anti-Hebbian long-term potentiation in the hippocampal feedback inhibitory circuit. Science, 315, 1262–6.

Lapointe, V., Morin, F., Ratte, S., Croce, A., Conquet, F. & Lacaille, J. C. 2004. Synapse-specific mGluR1-dependent long-term potentiation in interneurones regulates mouse hippocampal inhibition. J Physiol, 555, 125–35.

Leao, R. N., Mikulovic, S., Leao, K. E., Munguba, H., Gezelius, H., Enjin, A., Patra, K., Eriksson, A., Loew, L. M., Tort, A. B. & Kullander, K. 2012. OLM interneurons differentially modulate CA3 and entorhinal inputs to hippocampal CA1 neurons. Nat Neurosci, 15, 1524–30.

Lovett-Barron, M., Kaifosh, P., Kheirbek, M. A., Danielson, N., Zaremba, J. D., Reardon, T. R., Turi, G. F., Hen, R., Zemelman, B. V. & Losonczy, A. 2014. Dendritic inhibition in the hippocampus supports fear learning. Science, 343, 857–63.

Lovett-Barron, M., Turi, G. F., Kaifosh, P., Lee, P. H., Bolze, F., Sun, X. H., Nicoud, J. F., Zemelman, B. V., Sternson, S. M. & Losonczy, A. 2012. Regulation of neuronal input transformations by tunable dendritic inhibition. Nat Neurosci, 15, 423–30, S1–3.

Madisen, L., Zwingman, T. A., Sunkin, S. M., Oh, S. W., Zariwala, H. A., Gu, H., Ng, L. L., Palmiter, R. D., Hawrylycz, M. J., Jones, A. R., Lein, E. S. & Zeng, H. 2010. A robust and high-throughput Cre reporting and characterization system for the whole mouse brain. Nat Neurosci, 13, 133–40.

Mcbain, C. J. & Kauer, J. A. 2009. Presynaptic plasticity: targeted control of inhibitory networks. Curr Opin Neurobiol, 19, 254–62.

Mckay, B. M., Oh, M. M. & Disterhoft, J. F. 2013. Learning increases intrinsic excitability of hippocampal interneurons. J Neurosci, 33, 5499–506.

Muller, C. & Remy, S. 2014. Dendritic inhibition mediated by O-LM and bistratified interneurons in the hippocampus. Front Synaptic Neurosci, 6, 23.

Nabavi, S., Fox, R., Proulx, C. D., Lin, J. Y., Tsien, R. Y. & Malinow, R. 2014. Engineering a memory with LTD and LTP. Nature, 511, 348–52.

Nicholson, E. & Kullmann, D. M. 2021. Nicotinic receptor activation induces NMDA receptor independent long-term potentiation of glutamatergic signalling in hippocampal oriens interneurons. J Physiol, 599, 667–676.

O’Mara, S. M., Commins, S. & Anderson, M. 2000. Synaptic plasticity in the hippocampal area CA1-subiculum projection: implications for theories of memory. Hippocampus, 10, 447–56.

Oleskevich, S., Descarries, L. & Lacaille, J. C. 1989. Quantified distribution of the noradrenaline innervation in the hippocampus of adult rat. J Neurosci, 9, 3803–15.

Pelkey, K. A., Chittajallu, R., Craig, M. T., Tricoire, L., Wester, J. C. & Mcbain, C. J. 2017. Hippocampal GABAergic Inhibitory Interneurons. Physiol Rev, 97, 1619–1747.

Pelletier, J. G. & Lacaille, J. C. 2008. Long-term synaptic plasticity in hippocampal feedback inhibitory networks. Prog Brain Res, 169, 241–50.

Perez, Y., Morin, F. & Lacaille, J. C. 2001. A hebbian form of long-term potentiation dependent on mGluR1a in hippocampal inhibitory interneurons. Proc Natl Acad Sci U S A, 98, 9401–6.

Pouille, F. & Scanziani, M. 2004. Routing of spike series by dynamic circuits in the hippocampus. Nature, 429, 717–23.

Ran, I., Laplante, I., Bourgeois, C., Pepin, J., Lacaille, P., Costa-Mattioli, M., Pelletier, J., Sonenberg, N. & Lacaille, J. C. 2009. Persistent transcription- and translation-dependent long-term potentiation induced by mGluR1 in hippocampal interneurons. J Neurosci, 29, 5605–15.

Royer, S., Zemelman, B. V., Losonczy, A., Kim, J., Chance, F., Magee, J. C. & Buzsaki, G. 2012. Control of timing, rate and bursts of hippocampal place cells by dendritic and somatic inhibition. Nat Neurosci, 15, 769–75.

Sengupta, S., Peterson, T. R., Laplante, M., Oh, S. & Sabatini, D. M. 2010. mTORC1 controls fasting-induced ketogenesis and its modulation by ageing. Nature, 468, 1100–4.

Sharma, V., Sood, R., Khlaifia, A., Eslamizade, M. J., Hung, T. Y., Lou, D., Asgarihafshejani, A., Lalzar, M., Kiniry, S. J., Stokes, M. P., Cohen, N., Nelson, A. J., Abell, K., Possemato, A. P., Gal-Ben-Ari, S., Truong, V. T., Wang, P., Yiannakas, A., Saffarzadeh, F., Cuello, A. C., Nader, K., Kaufman, R. J., Costa-Mattioli, M., Baranov, P. V., Quintana, A., Sanz, E., Khoutorsky, A., Lacaille, J. C., Rosenblum, K. & Sonenberg, N. 2020. eIF2alpha controls memory consolidation via excitatory and somatostatin neurons. Nature, 586, 412–416.

Sun, Y., Nguyen, A. Q., Nguyen, J. P., Le, L., Saur, D., Choi, J., Callaway, E. M. & Xu, X. 2014. Cell-type-specific circuit connectivity of hippocampal CA1 revealed through Cre-dependent rabies tracing. Cell Rep, 7, 269–80.

Szonyi, A., Sos, K. E., Nyilas, R., Schlingloff, D., Domonkos, A., Takacs, V. T., Posfai, B., Hegedus, P., Priestley, J. B., Gundlach, A. L., Gulyas, A. I., Varga, V., Losonczy, A., Freund, T. F. & Nyiri, G. 2019. Brainstem nucleus incertus controls contextual memory formation. Science, 364.

Taniguchi, H., He, M., Wu, P., Kim, S., Paik, R., Sugino, K., Kvitsiani, D., Fu, Y., Lu, J., Lin, Y., Miyoshi, G., Shima, Y., Fishell, G., Nelson, S. B. & Huang, Z. J. 2011. A resource of Cre driver lines for genetic targeting of GABAergic neurons in cerebral cortex. Neuron, 71, 995–1013.

Taube, J. S. 1993. Electrophysiological properties of neurons in the rat subiculum in vitro. Exp Brain Res, 96, 304–18.

Topolnik, L., Azzi, M., Morin, F., Kougioumoutzakis, A. & Lacaille, J. C. 2006. mGluR1/5 subtype-specific calcium signalling and induction of long-term potentiation in rat hippocampal oriens/alveus interneurones. J Physiol, 575, 115–31.

Tyan, L., Chamberland, S., Magnin, E., Camire, O., Francavilla, R., David, L. S., Deisseroth, K. & Topolnik, L. 2014. Dendritic inhibition provided by interneuron-specific cells controls the firing rate and timing of the hippocampal feedback inhibitory circuitry. J Neurosci, 34, 4534–47.

Udakis, M., Pedrosa, V., Chamberlain, S. E. L., Clopath, C. & Mellor, J. R. 2020. Interneuron-specific plasticity at parvalbumin and somatostatin inhibitory synapses onto CA1 pyramidal neurons shapes hippocampal output. Nat Commun, 11, 4395.

Van Hooft, J. A., Giuffrida, R., Blatow, M. & Monyer, H. 2000. Differential expression of group I metabotropic glutamate receptors in functionally distinct hippocampal interneurons. J Neurosci, 20, 3544–51.

Vasuta, C., Artinian, J., Laplante, I., Hebert-Seropian, S., Elayoubi, K. & Lacaille, J. C. 2015. Metaplastic Regulation of CA1 Schaffer Collateral Pathway Plasticity by Hebbian MGluR1a-Mediated Plasticity at Excitatory Synapses onto Somatostatin-Expressing Interneurons. eNeuro, 2.

Whitlock, J. R., Heynen, A. J., Shuler, M. G. & Bear, M. F. 2006. Learning induces long-term potentiation in the hippocampus. Science, 313, 1093–7.

Younts, T. J., Monday, H. R., Dudok, B., Klein, M. E., Jordan, B. A., Katona, I. & Castillo, P. E. 2016. Presynaptic Protein Synthesis Is Required for Long-Term Plasticity of GABA Release. Neuron, 92, 479–492.

Yuan, S. & Burrell, B. D. 2013. Endocannabinoid-dependent long-term depression in a nociceptive synapse requires coordinated presynaptic and postsynaptic transcription and translation. J Neurosci, 33, 4349–58.

