## Supplementary figures and table for "Long-term potentiation at pyramidal cell to somatostatin interneuron synapses controls hippocampal network plasticity and memory"

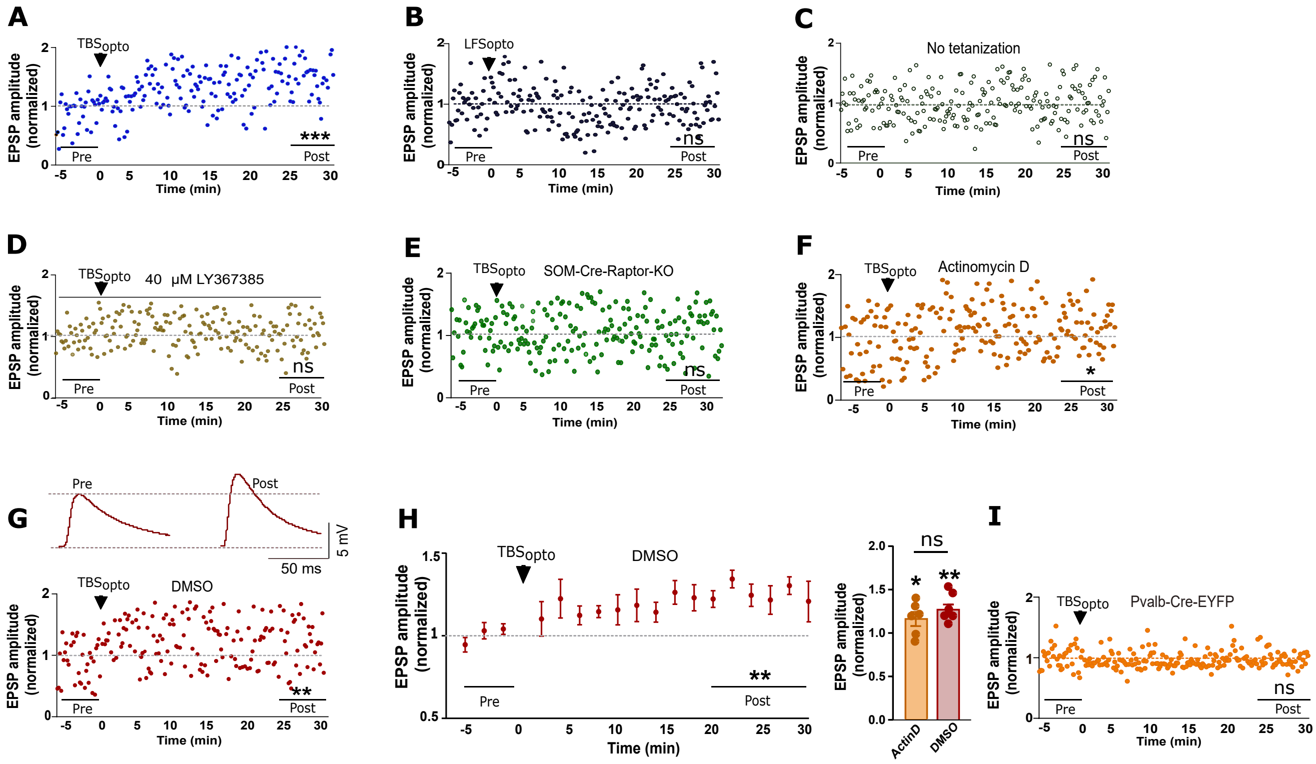
A
TBSopto
EPSP amplitude (normalized)
Pre
Post
***
Time (min)
B
LFSopto
ns
C
No tetanization
ns
D
40 µM LY367385
ns
E
SOM-Cre-Raptor-KO
ns
F
Actinomycin D
*
G
50 ms
5 mV
DMSO
**
H
ActinD
DMSO
I
Pvalb-Cre-EYFP
ns

**Figure S1. Optogenetically-induced LTP at PC-SOM synapses in representative cells, related to Figure 1.**
(A to E) Time plots of light-evoked EPSP amplitude from representative SOM cells (EPSPs shown in Figure 1B to 1G) showing LTP at PC-SOM synapses after $TBS_{opto}$ (A), but not after $LFS_{opto}$ (B), no tetanization (C), $TBS_{opto}$ in LY367385 (D), or $TBS_{opto}$ in SOM-Cre-Raptor-KO mice (E). Paired t-tests at 25-30 min after induction (Post) relative to baseline (Pre), *** $p < 0.0001$, ns $p > 0.05$.
(F and G) Time plots of light-evoked EPSP amplitude from representative SOM cells (EPSPs shown in Figure 1G for cell in F), showing LTP at PC-SOM synapses after $TBS_{opto}$ after incubation with the transcription inhibitor actinomycin D (F) or its vehicle DMSO (G; EPSPs above). Paired t-tests, * $p < 0.05$, ** $p < 0.01$.
(H) Left: time plot of light-evoked EPSP amplitude for all SOM cells tested showing LTP at PC-SOM synapses following $TBS_{opto}$ after incubation with DMSO (n = 6 cells). Right: Summary graph of EPSP amplitude for all cells at 20-30 min post induction, showing similar LTP after $TBS_{opto}$ after incubation in actinomycin D or its vehicle DMSO. Paired t-tests (Pre vs Post) and unpaired t-test (Post Actin D vs Post DMSO), * $p < 0.05$, ** $p < 0.01$, ns $p > 0.05$.
(I) Time plots of light-evoked EPSP amplitude from representative Pvalb cell (EPSPs shown in Figure 1I) showing absence of LTP at PC-Pvalb synapses after $TBS_{opto}$. Paired t-test, ns $p > 0.05$.

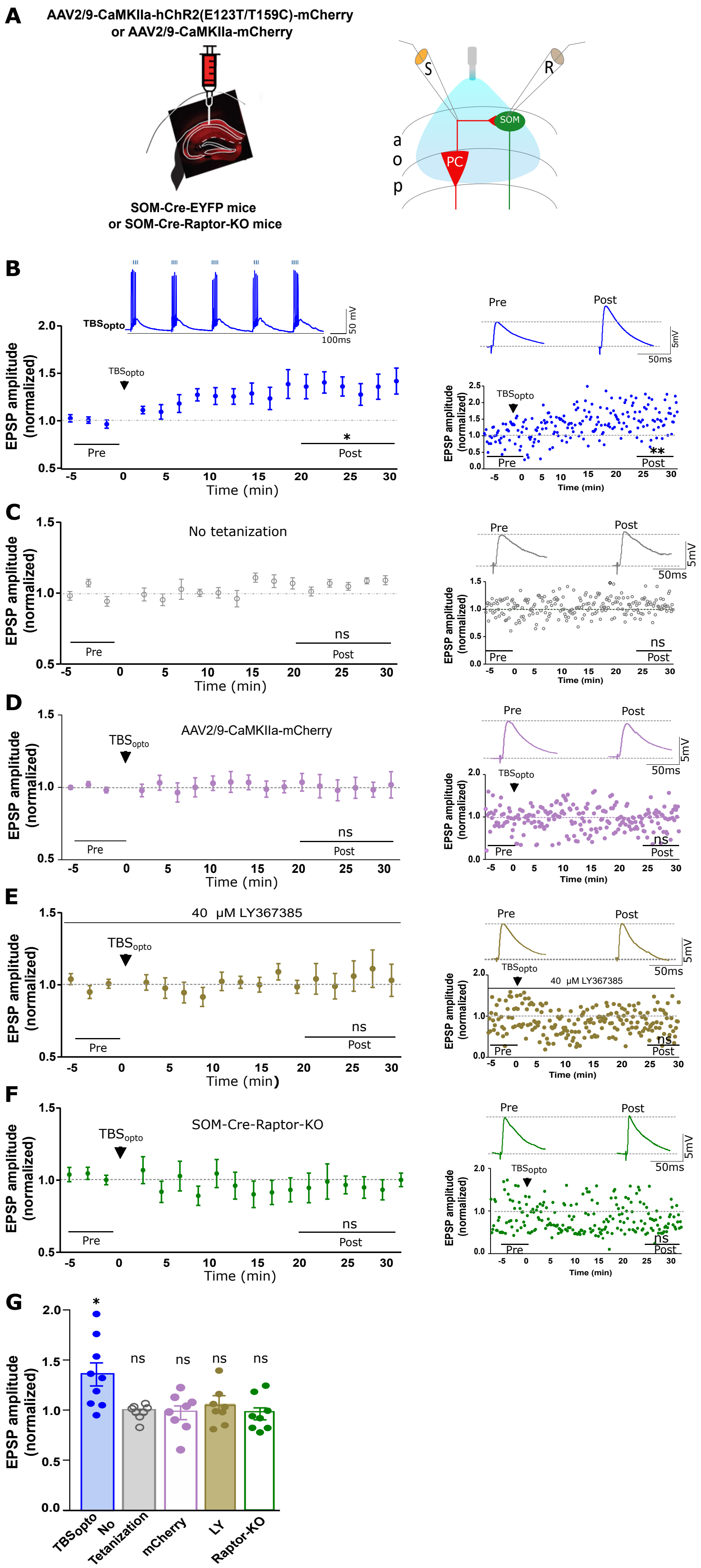
A
AAV2/9-CaMKIIa-hChR2(E123T/T159C)-mCherry
or AAV2/9-CaMKIIa-mCherry
SOM-Cre-EYFP mice
or SOM-Cre-Raptor-KO mice
S
R
SOM
PC
a
o
p
B
TBSopto
50 mV
100ms
EPSP amplitude (normalized)
Time (min)
Pre
Post
*
**
5mV
50ms
C
No tetanization
ns
D
AAV2/9-CaMKIIa-mCherry
E
40 μM LY367385
F
SOM-Cre-Raptor-KO
G
TBSopto
No Tetanization
mCherry
LY
Raptor-KO

**Figure S2. Optogenetically-induced LTP of electrically-evoked EPSPs in SOM cells, related to Figure 1.**

(A) Left: schematic of experimental paradigm with viral injections in dorsal hippocampus of SOM-Cre-EYFP or SOM-Cre-Raptor-KO mice. Right: diagram of local optogenetic stimulation of CA1 pyramidal cells, electrical stimulation of afferents and whole cell recording from SOM cells. (B) Left: time plot of electrically-evoked EPSP amplitude for all SOM cells, showing LTP at 20-30 min following $TBS_{opto}$ (n = 9 cells). Inset above: EPSP summation and cell firing during $TBS_{opto}$. Right: example of EPSPs (top) and time plot (bottom) from a representative cell, before (Pre) and 25-30 min after (Post) induction. Paired t-tests; ** $p < 0.01$, * $p < 0.05$.

(C - F) Time plots of electrically-evoked EPSP amplitude for all SOM cells (left) and example of EPSPs (top) and time plot (bottom) from a representative cell (right), showing absence of LTP of electrically-evoked EPSP amplitude following no tetanization (C; n = 8 cells), or following $TBS_{opto}$ in slices from mice with control CaMKIIa-mCherry injection without hChR2 (D; n = 8 cells), $TBS_{opto}$ in the presence of the mGluR1a antagonist LY367385 (40 µM) (E; n = 8 cells), or $TBS_{opto}$ in SOM-Cre-Raptor-KO mice (F; n = 8 cells). Paired t-tests; ns, $p > 0.05$.

(G) Summary graph of EPSP amplitude for all cells at 20-30 min post induction, showing LTP after $TBS_{opto}$, but no LTP in absence of tetanization, or after $TBS_{opto}$ in mice injected with mCherry, in LY367385, or in SOM-Cre-Raptor-KO mice. Paired t-tests (Pre vs Post); * $p < 0.05$, ns $p > 0.05$.

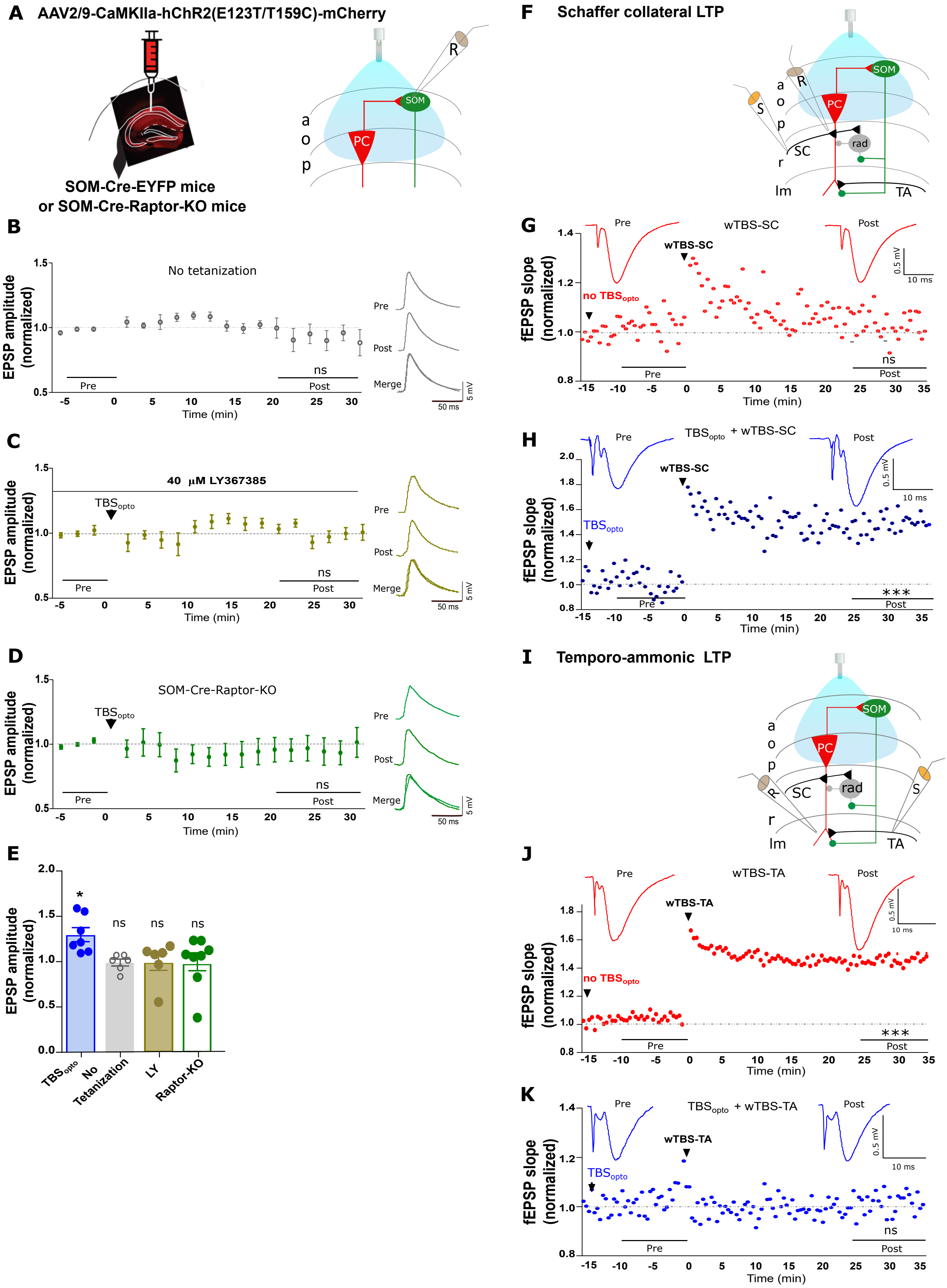
A AAV2/9-CaMKIIa-hChR2(E123T/T159C)-mCherry
SOM-Cre-EYFP mice
or SOM-Cre-Raptor-KO mice
B No tetanization
EPSP amplitude (normalized)
Time (min)
Pre
Post
Merge
C 40 μM LY367385
TBSopto
D SOM-Cre-Raptor-KO
E
TBSopto
No Tetanization
LY
Raptor-KO
F Schaffer collateral LTP
G wTBS-SC
no TBSopto
fEPSP slope (normalized)
H TBSopto + wTBS-SC
I Temporo-ammonic LTP
J wTBS-TA
no TBSopto
K TBSopto + wTBS-TA

**Figure S3. Whole-field $TBS_{opto}$-induced LTP at PC-SOM synapses and differential regulation of LTP at SC-PC and TA-PC synapses, related to Figure 2.**
(A) Left: schematic of experimental paradigm with viral injections in dorsal hippocampus of SOM-Cre-EYFP or SOM-Cre-Raptor-KO mice. Right: diagram of whole-field optogenetic stimulation of CA1 pyramidal cells and whole cell recording from SOM interneurons.
(B-D) Left: time plots of light-evoked EPSP amplitude for all SOM cells, showing absence of LTP at PC-SOM synapses following no tetanization (B; n = 6 cells), $TBS_{opto}$ in the presence of the mGluR1a antagonist LY367385 (C; n = 6 cells), or $TBS_{opto}$ in SOM-Cre-Raptor-KO mice (D; n =8 cells). Right: representative example of average EPSPs before (Pre) and 20-30 min after (Post) induction in each treatment condition. Paired t-tests, ns $p > 0.05$.
(E) Summary graph of light-evoked EPSP amplitude for all cells at 20-30 min post induction, showing LTP after $TBS_{opto}$, but not in absence of tetanization, nor after $TBS_{opto}$ in LY367385 or SOM-Cre-Raptor-KO mice. Paired t-tests; * $p < 0.05$, ns $p > 0.05$.
(F) Diagram of whole-field optogenetic stimulation of CA1 pyramidal cells, with electrical stimulation and recording of SC-PC fEPSPs in stratum radiatum.
(G-H) Time plots of fEPSP slope (with average fEPSPs above) from representative slices, showing that LTP at SC-PC synapses (G; red) is facilitated by prior application of $TBS_{opto}$ (H; blue). Paired t-tests, *** $p < 0.001$, ns, $p > 0.05$.
(I) Diagram of whole-field optogenetic stimulation of CA1 pyramidal cells, with electrical stimulation and recording of TA-PC fEPSPs in stratum lacunosum-moleculare.
(J-K) Time plots of fEPSP slope (with average fEPSPs above) from representative slices, showing that LTP at TA-PC synapses (J; red) is depressed by prior application of $TBS_{opto}$ (H; blue). Paired t-tests, *** $p < 0.001$, ns, $p > 0.05$.

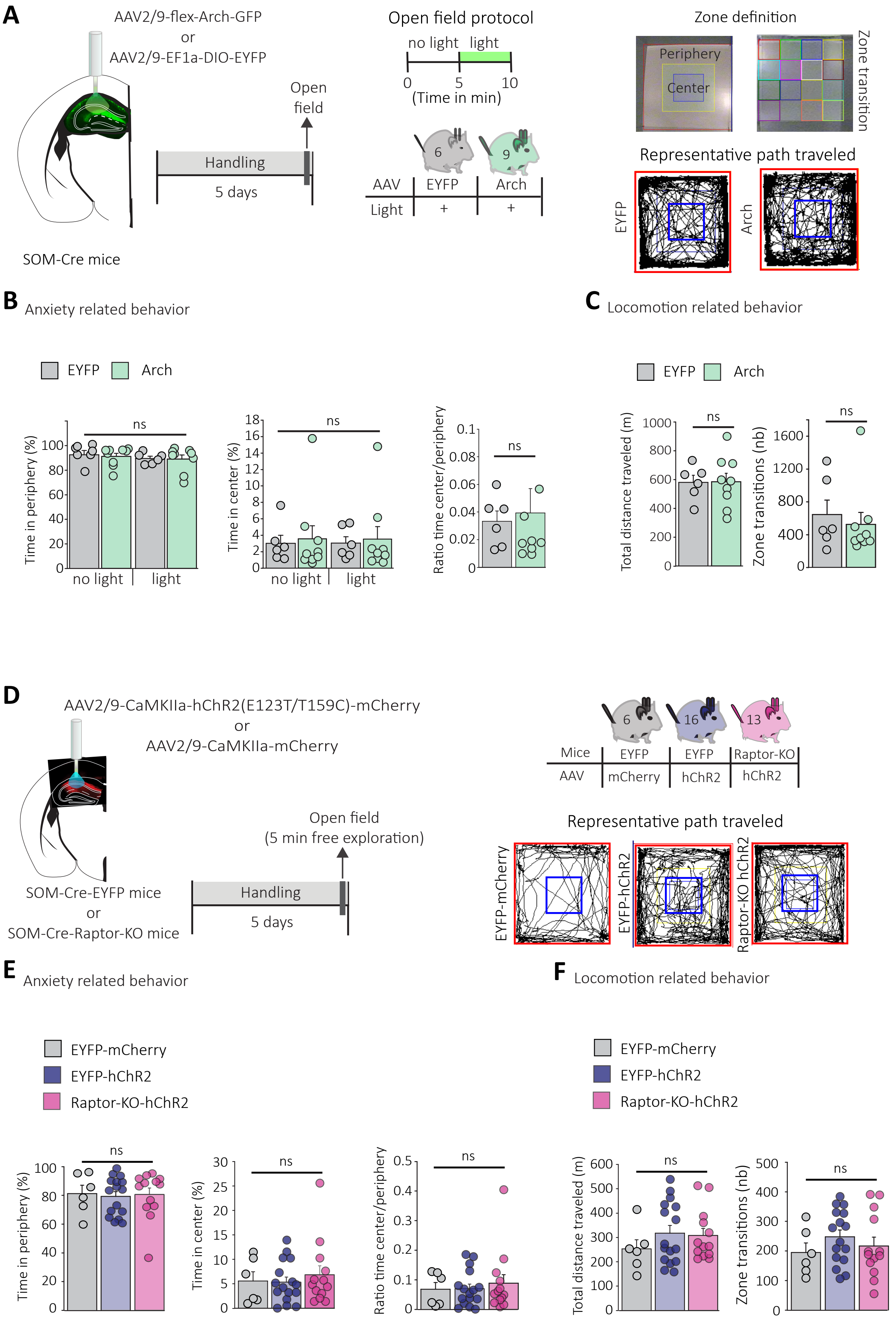
A
AAV2/9-flex-Arch-GFP
or
AAV2/9-EF1a-DIO-EYFP
SOM-Cre mice
Handling
5 days
Open field
Open field protocol
no light
light
0
5
10
(Time in min)
6
9
AAV
EYFP
Arch
Light
+
+
Zone definition
Periphery
Center
Zone transition
Representative path traveled
EYFP
Arch
B Anxiety related behavior
EYFP
Arch
Time in periphery (%)
ns
no light
light
Time in center (%)
ns
no light
light
Ratio time center/periphery
ns
C Locomotion related behavior
EYFP
Arch
Total distance traveled (m)
ns
Zone transitions (nb)
ns
D
AAV2/9-CaMKIIa-hChR2(E123T/T159C)-mCherry
or
AAV2/9-CaMKIIa-mCherry
SOM-Cre-EYFP mice
or
SOM-Cre-Raptor-KO mice
Handling
5 days
Open field
(5 min free exploration)
6
16
13
Mice
EYFP
EYFP
Raptor-KO
AAV
mCherry
hChR2
hChR2
Representative path traveled
EYFP-mCherry
EYFP-hChR2
Raptor-KO hChR2
E Anxiety related behavior
EYFP-mCherry
EYFP-hChR2
Raptor-KO-hChR2
Time in periphery (%)
ns
Time in center (%)
ns
Ratio time center/periphery
ns
F Locomotion related behavior
EYFP-mCherry
EYFP-hChR2
Raptor-KO-hChR2
Total distance traveled (m)
ns
Zone transitions (nb)
ns

**Figure S4. Normal anxiety and locomotion in open field tests, related to Figure 3.**
(A) Left: schematic of experimental paradigm with injections of AAV2/9-flex-Arch-GFP or AAV2/9-EF1a-DIO-EYFP in dorsal CA1 hippocampus of SOM-Cre mice. Middle: diagram of protocol during two open field test episodes, first without light and second with light for optogenetic silencing of SOM interneurons. Right: (top) zone separations used for analysis and (bottom) representative examples of path traveled by mice with EYFP (left) or Arch (right) expression in open field test. (B) Summary graphs for all mice showing unchanged time spent in periphery (left) and center (middle), and ratio of time in center/periphery (right), of mice that received light activation of Arch (Arch; n=9 mice, green) relative to mice with light stimulation without Arch (EYFP; n=6 mice, grey) during open field tests without and with light, indicating normal anxiety level in both groups. Two-way repeated measures ANOVA (time spent in periphery and center); Mann-Whitney Rank Sum Test (ratio of time center/periphery); ns p>0.05.
(C) Summary graphs for all mice showing unchanged total distance traveled and zone transitions in mice that received light activation of Arch (green) relative to mice with light stimulation without Arch (gray) during open field tests, indicating normal locomotion in both groups. Unpaired t-test (distance traveled) and Mann-Whitney Rank Sum Test (zone transitions), ns p>0.05.
(D) Left: Schematic of experimental paradigm with injections of AAV2/9-CaMKIIa-hChR2(E123T/T159C)-mCherry or AAV2/9-CaMKIIa-mCherry in dorsal CA1 hippocampus of SOM-Cre-EYFP and SOM-Cre-Raptor-KO mice, and diagram of behavioral testing sequence (open field test). Right: representative examples of path traveled in open field test by SOM-Cre-EYFP mice with mCherry or hChR2 expression, and SOM-Cre-Raptor-KO mice with hChR2 expression.
(E) Summary graphs for all mice showing unchanged time spent in periphery (left) and center (middle), and ratio of time in center/periphery (right), of SOM-Cre-EYFP mice with mCherry expression (n=6 mice, grey) or hChR2 expression (n=16 mice, violet), and SOM-Cre-Raptor-KO mice with hChR2 expression (n=13 mice, pink), indicating normal anxiety level. Kruskal-Wallis one-way ANOVA on Ranks, ns p>0.05. (F) Summary graphs for all mice showing unchanged total distance traveled and zone transition number of SOM-Cre-EYFP mice with mCherry expression (n=6 mice, pink) or hChR2 expression (n=15 mice, blue), and SOM-Cre-Raptor-KO mice with hChR2 expression (n=13 mice, green), indicating normal locomotion. Kruskal-Wallis one-way ANOVA on Ranks (distance traveled), one-way ANOVA (zone transitions), ns p>0.05.

**Table S2. Light-evoked EPSP and electrical-evoked EPSP amplitude pre-TBS in all related experiments, related to Figures 1, 4, S2, and S3.**

| **Experiments Figure 1** | **N** | **EPSP Amplitude (mV) Mean±SEM** | **Statistic** |
|---|---|---|---|
| $TBS_{opto}$ | 17 | 7.786±0.6659 | Dunn's multiple comparisons test |
| $LFS_{opto}$ | 8 | 7.106±0.6369 | P= 0.9999 |
| No tetanization | 13 | 7.811±0.9687 | P= 0.9999 |
| LY | 10 | 8.533±0.6725 | P= 0.9999 |
| Raptor-KO | 15 | 6.448±0.4820 | P= 0.7334 |
| ActinD | 6 | 5.840±0.8812 | P= 0.6247 |
| DMSO | 6 | 6.106±0.6708 | P= 0.9999 |
| Pvalb | 8 | 4.872±0.4452 | P= 0.0167* |
| **Experiments Figure S2** | **N** | **EPSP Amplitude (mV) Mean±SEM** | **Statistic** |
| $TBS_{opto}$ | 9 | 4.339±0.4648 | Dunn's multiple comparisons test |
| No tetanization | 8 | 5.967±0.5759 | P= 0.2202 |
| mCherry | 8 | 6.026±0.7012 | P= 0.2202 |
| LY | 8 | 6.270±0.5736 | P= 0.0563 |
| Raptor-KO | 8 | 5.631±0.6788 | P= 0.7228 |
| **Experiments Figure S3** | **N** | **EPSP Amplitude (mV) Mean±SEM** | **Statistic** |
| $TBS_{opto}$ | 7 | 8.852±0.5967 | Dunn's multiple comparisons test |
| No tetanization | 6 | 7.375±1.299 | P= 0.9999 |
| LY | 6 | 10.52±0.4438 | P= 0.4059 |
| Raptor-KO | 8 | 6.532±0.7446 | P= 0.2394 |
| **Experiments Figure 4** | **N** | **EPSP Amplitude (mV) Mean±SEM** | **Statistic** |
| B | 8 | 4.540±0.4835 | 0.2089 |
| C | 7 | 4.248±0.3853 | 0.0937 |
| D | 8 | 6.427± 0.4251 | 0.5308 |
| E | 8 | 5.367±0.3607 | 0.9694 |
| F | 10 | 5.660±0.4683 | F (4, 36) = 3.802 |
