## Supplementary material for "Long-term potentiation at pyramidal cell to somatostatin interneuron synapses controls hippocampal network plasticity and memory": Star methods

### KEY RESOURCES TABLE

| REAGENT or RESOURCE | SOURCE | IDENTIFIER |
|---|---|---|
| **Antibodies** | | |
| Rabbit polyclonal anti-GFP | ThermoFisher Scientific | Cat# A11122; RRID : AB_221569 |
| Alexa Fluor 594-conjugated goat anti-rabbit IgG | Jackson Immunoresearch Laboratories | Cat# 111-585-003; RRID : AB_2338059 |
| **Chemicals, Peptides, and Recombinant Proteins** | | |
| LY367385 | Tocris Bioscience | Cat# 1237; CAS# 198419-91-9 |
| Actinomycin D | Sigma | Cat# 114666; CAS# 50-76-0 |
| DL-2-Amino-5-phosphonopentanoic acid (DL-AP5) | Abcam | Cat# ab120004 CAS# 76326-31-3 |
| SR-95531 (Gabazine) | Abcam | Cat# ab120042 CAS# 104104-50-9 |
| **Experimental Models: Organisms/Strains** | | |
| Mouse: $Sst^{ires\text{-}Cre}$: $Sst^{tm2.1(cre)Zjh}$/J | The Jackson Laboratory (Taniguchi et al., 2011) | JAX: 013044 |
| Mouse: $Rosa26^{lsl\text{-}EYFP}$: B6.Cg-*Gt(ROSA)26Sor*$^{tm3(CAG\text{-}EYFP)Hze}$/J | The Jackson Laboratory (Madisen et al., 2010) | JAX: 007903 |
| Mouse: $Rptor^{fl/fl}$: B6.Cg-$Rptor^{tm1.1Dmsa}$/J | The Jackson Laboratory (Sengupta et al., 2010) | JAX: 013188 |
| Mouse: $Sst^{ires\text{-}Cre}$;$Rosa26^{lsl\text{-}EYFP}$;$Rptor^{fl/fl}$ | (Artinian et al., 2019) | N/A |
| Mouse: $Pvalb^{ires\text{-}Cre}$: B6;129P2-$Pvalb^{tm1(cre)Arbr}$/J | The Jackson Laboratory (Hippenmeyer et al., 2005) | JAX: 008069 |
| **Recombinant DNA** | | |
| AAV2/9-CaMKIIa-hChR2(E123T/T159C)-mCherry | UPenn Vector Core; Canadian Neurophotonics Platform | Addgene 35512 |
| AAV2/9-CaMKIIa-mCherry | UPenn Vector Core; Canadian Neurophotonics Platform | Addgene 114469 |
| AAV2/9-EF1a-DIO-EYFP | UPenn Vector Core; Canadian Neurophotonics Platform | Addgene 27056 |
| AAV2/9-flex-Arch-GFP | UPenn Vector Core | Addgene 22222 |
| **Software and Algorithms** | | |
| pClamp 10.5, 10.7 | Molecular Devices | N/A |
| Clampfit 10.5, 10.7 | Molecular Devices | N/A |
| Multiclamp | Molecular Devices | N/A |
| PolyLite | Mightex | N/A |
| Polyscan2 | Mightex | N/A |

| | | |
|---|---|---|
| SimplePCI | Compix Inc, Imaging Sytems | N/A |
| FreezeFrame Actimetrics | Coulbourn Instruments | N/A |
| Smart 3.0 tracking system | Panlab | N/A |
| Graph Pad Prism 6 | GraphPad | N/A |
| SigmaPlot 11 | Systat Software Inc | N/A |
| Photoshop CS3 | Adobe | N/A |
| Illustrator 17.1 | Adobe | N/A |
